# CSFeatures improves identification of cell type-specific differential features in single-cell and spatial omics data

**DOI:** 10.1101/2025.05.21.655244

**Authors:** Rufeng Li, Yongkang Li, Heyang Hua, Zhen Li, Yu Zhang, Yufei Yao, Shengquan Chen, Yungang Xu

## Abstract

Recent advancements in single-cell and spatial omics have enabled the analysis of gene expression patterns in cells and tissues with unprecedented precision. Genes that exhibit significant variation across different cell types or states are typically closely linked to cellular functional states and disease processes. However, current differential analysis methods often fail to adequately account for cell type specificity when identifying differentially expressed genes. CSFeatures effectively addresses this issue by comprehensively considering the expression level, the smoothness of expression distribution, and the proportion of gene expression in the target cell population as well as in each of the other cell populations. Through comprehensive experiments on eight single-cell RNA-seq datasets, it was demonstrated that CSFeatures can effectively identify cell type-specific genes that are highly expressed in the target cell population while being lowly expressed in all other populations. Moreover, it was validated that the underlying principle of CSFeatures can be generalized to other omics data, including single-cell ATAC-seq, spatial transcriptomics, and spatial ATAC-seq data. Furthermore, the differential features identified by CSFeatures are enriched in pathways, functional categories, and regulatory regions that are highly relevant to the specific functional states of the target cell population, providing insights into its gene regulation.

## Introduction

Single-cell and spatial omics analysis has enabled scientists to understand cellular heterogeneity with unprecedented detail, providing profound insights into the molecular processes driving cell differentiation and complex diseases(1–3). In the analysis of single-cell RNA sequencing (scRNA-seq) data, identifying genes that exhibit significant differences between cell populations or states is a critical step following cell clustering. These differentially expressed genes (DEGs) help elucidate the distinct functional states and biological roles of different cell populations, offering valuable insights into mechanisms underlying diseases(4–6). However, technical noise often masks molecular variations between cells, making it challenging to identify cell type-specific DEGs, which are highly expressed in the target cell population while showing extremely weak expression in others.

Currently, commonly used differential analysis (DA) methods for scRNA-seq data include the Wilcoxon rank-sum test, Student’s t-test, MAST(7), among others(8). Recently, Pullin and colleagues conducted a benchmark study on DA methods for scRNA-seq data, highlighting the effectiveness of simple methods such as the Wilcoxon rank-sum test, Student’s t-test, and logistic regression by evaluating their ability to recover expert-annotated marker genes(8). These methods are often integrated into analysis frameworks such as Seurat(9) and Scanpy(10). Despite their widespread use, these mainstream tools typically combine all non-target cell populations into a single “other” group for gene expression comparisons with the target population. However, imbalanced sample sizes and substantial biological heterogeneity within the ‘other’ group present challenges for differential analysis. Specifically, many of the top-ranked genes identified by commonly used DA methods lack expected cell type specificity. For instance, some highly ranked genes may be strongly expressed in the target population but also show substantial expression in certain other populations. Conversely, certain genes that are weakly expressed in other populations may not exhibit strong expression in the target population either. Moreover, even when comparing differential gene expression between the two cell populations, these challenges may also arise. In subsequent sections, we provide a thorough demonstration and discussion of these limitations.

Moreover, similar analytical challenges are encountered in the analysis of single-cell assay for transposase-accessible chromatin sequencing (scATAC-seq), spatial transcriptomics (ST), and spatial ATAC-seq (spaATAC-seq) data. scATAC-seq data are typically sparse and often represented as binary matrices. Currently, differential region identification is commonly performed using logistic regression (LR), which is better suited for handling binary response variables(11). However, as with scRNA-seq data, the differentially accessible regions identified by LR for scATAC-seq data frequently lack cell type specificity. In some cases, even the top-ranked regions show low chromatin accessibility in the target cell population. We subsequently confirmed this across various datasets. Likewise, in the analysis of spatial omics data (including both ST and spaATAC-seq data), after identifying spatial domains, it becomes crucial to link these domains to biological functions by identifying differential features specifically associated with different domains(12,13). Unfortunately, the features detected by current DA methods do not always display clear spatial patterns(13–16), which complicates their use in precise downstream analyses, such as constructing cell-cell communication networks or mapping developmental trajectories. Overall, current methods for identifying differential features between cell populations in single-cell and spatial omics often fail to capture cell type or spatial domain specificity. These limitations primarily arise because existing approaches do not comprehensively account for feature activity across all cell populations. Moreover, these limitations introduce biases in downstream analyses and poses challenges for biologists when selecting key features for further validation. For simplicity, we will refer to both cell type and spatial domain specificity collectively as cell type specificity throughout the remainder of this paper.

In this work, we first demonstrated that the lack of cell type specificity in identifying differential features is a common issue in current DA methods via multiple examples. To address this limitation, we propose CSFeatures, a method for identifying cell type-specific differential features in single-cell and spatial omics data. Taking scRNA-seq data as an example, for each gene, CSFeatures comprehensively considers its average expression level, the smoothness of its expression distribution, and the proportion of cells expressing the gene across different cell populations. We validated CSFeatures on eight scRNA-seq datasets generated from different technologies, including the 10x Genomics Chromium, Drop-seq(^17^), and CEL-seq2(^18^). Compared to commonly used DA methods, the top-ranked genes identified by CSFeatures exhibit high expression in the target cell population and extremely low expression in other populations. Furthermore, we found that the underlying principle of CSFeatures can be generalized to other omics data. With slight adjustments, the method performs equally well on scATAC-seq data, effectively identifying cell type-specific differentially accessible regions. Additionally, CSFeatures can leverage appropriate embedding methods to integrate spatial coordinates, enabling the identification of features with spatial patterns in spatial omics data. Moreover, the differential features identified by CSFeatures are closely aligned with the specialized functions of the target cell population. This capability is essential for uncovering specialized cellular functions and elucidating complex biological mechanisms.

## Methods

### Data preprocessing

All datasets used in this study are publicly available, with some providing preprocessed data that have undergone normalization. For these preprocessed datasets, we directly proceeded with differential analyses. For datasets provided as raw count matrices, we applied consistent preprocessing methods tailored to each omics data. Specifically, for single-cell RNA sequencing (scRNA-seq) and spatial transcriptomics (ST) data, we filtered out genes expressed in fewer than three cells or spots and then normalized the data using the *NormalizeData* function in Seurat(9) v5.0.3. For single-cell assay for transposase-accessible chromatin sequencing (scATAC-seq) and spatial ATAC-seq (spaATAC-seq) data, we filtered out regions present in less than 1% of the cells or spots(19), followed by reweighting region occurrence frequencies using term frequency-inverse document frequency (TF-IDF) transformation(20,21).

### The model of CSFeatures

For each gene, CSFeatures comprehensively considers its average expression level, the smoothness of its expression distribution, and the proportion of cells expressing the gene within each cell population. We devised the expression index (EI) metric to quantify the degree of cell type specificity for each gene. The underlying principle of CSFeatures is broadly applicable and can be generalized to various omics data, including scRNA-seq, scATAC-seq, ST, and spaATAC-seq data.

Taking scRNA-seq data as an example, the input of CSFeatures consists of a preprocessed gene expression matrix **X**, with the dimension of *n* × *g*, where *n* represents the number of cells and *g* represents the number of genes. Additionally, **Y** is a vector that contains the labels to be investigated, such as cell type labels or clustering labels, for the *n* cells. For clarity in explaining the subsequent calculation process, we take the gene *k* in the target cell population as an example.

Firstly, to quantify the relationships between cells, we construct a cell similarity matrix **C** with the dimension of *n* × *n*, where the **C***_ij_* represents the similarity between cell *i* and cell *j*. We perform principal component analysis (PCA) on the preprocessed expression matrix **X** to reduce its dimensionality to 50 by default, resulting in a dimension-reduced matrix **X**_pca_. Next, the distance matrix **D**, which measures the overall similarity of gene expression patterns between cells, is computed based on **X**_pca_ using Euclidean distance. **C** can be derived using the following formula:

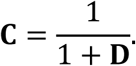

Secondly, to reflect the expression level of the gene *k*, we calculate the mean expression *M_tk_* within the target cell population. Assume that the target population consists of *p* cells. The calculation formula for *M_tk_* is as follows:

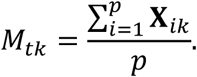

Third, to assess the variability of gene expression across cells, we construct a graph for the target cell population based on *C*, where the expression levels of gene *k* are represented as vertices, and the similarities between cells are represented as edge weights. The gene expression variability, *S_tk_*, is then quantified by calculating the smoothness of this graph, which is defined as

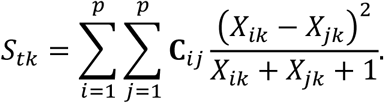

Moreover, to account for the fact that cell type-specific genes are expected to be highly expressed within the target cell population while displaying low expression in other cells, we compute the local maximum mean *L_ok_* for gene *k*. We define the adjacency matrix **A**, which reflects the *q* nearest neighbors of each cell based on **D**, where the default value of *q* is 20. For cell *i* outside the target cell population, the average expression of gene *k* across the *q* nearest neighboring cells is calculated. *L_ok_* represents the maximum of these values. The indices of the *q* nearest neighboring cells for the cell *i* are denoted by α_*i*1_, α*_i2_*, …, α_*i,q*-1_, α*_iq_*, respectively. The formula for calculating *L_ok_* is as follows:

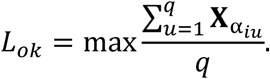

We then integrate *M_tk_*, *S_tk_* and *L_ok_* to define *V_k_*, the expression term that evaluates the expression level of the gene *k*. *V_k_* is calculated as

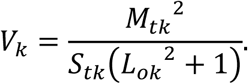

Furthermore, we expect that genes with cell type specificity will exhibit high expression in the majority of cells within the target cell population, while showing minimal or no expression in other cell populations. To further refine this distinction, we also take into account the proportion of cells with non-zero expression levels. Let *P_tk_* represent the expression proportion of the gene *k* within the target cell population, and *P_rk_* represent its expression proportion in the *r* -th population outside the target cell population. Assume there are ℎ cell populations excluding the target population. The proportion term *P_k_* is defined as follows:

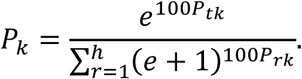

Finally, we integrate the expression term *V_k_* and proportion term *P_k_* to compute the expression index (EI) for gene *k*. A higher EI value indicates stronger cell type specificity for the gene. *El_k_* is calculated as

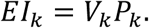

The process outlined above describes in detail how CSFeatures identifies cell type-specific differentially expressed genes in scRNA-seq data. For other omics data, there are some variations. In scATAC-seq data, when identifying cell type-specific differentially accessible regions, the main difference lies in the data preprocessing, which employs a different approach compared to scRNA-seq data. For ST data, spatial coordinates are integrated with gene expression during the dimensionality reduction step using SpaGCN(13), replacing PCA (Supplementary Text 1). For spaATAC-seq data, in addition to using TF-IDF during preprocessing, spaPeakVAE(22) is applied to integrate spatial coordinates with chromatin accessibility data for cell embedding.

### The implementation of the baseline methods

We implemented commonly used differential analysis methods by the *FindMarkers* function in Seurat v5.0.3, applying various test methods by adjusting the *test.use* parameter:

wilcox: The Wilcoxon rank-sum test, as the default non-parametric test method in the *FindMarkers* function, is widely used in differential analysis for its robustness to non-normal distributions.

bimod(23): The likelihood-ratio test is used for fitting a two-component mixture model, which is effective in capturing expression level changes.

t: The Student’s t-test is appropriate for normally distributed data but is sensitive to outliers.

negbinom: The negative binomial test is used for overdispersed count data, which is typical in scRNA-seq data.

poisson: The Poisson test is used for count data, though it is limited by overdispersion commonly seen in single-cell omics data.

LR: The logistic regression is used to model binary outcomes and assess the likelihood of gene expression being associated with specific conditions.

MAST(7): A hurdle model designed for single-cell data accounts for both zero-inflation and overdispersion, but it is computationally intensive.

### Simulation of scRNA-seq data

We simulated scRNA-seq data using splatter(24) v1.28.0, generating a dataset consisting of 2,000 cells and 8,000 genes, partitioned into five distinct clusters. Cluster 0 was defined as the target population, and the expression patterns of specific genes were modified. Specifically, 200 genes were set to be expressed in 50% of cells in cluster 0, while being expressed in only 2% of cells in the other clusters. Additionally, another set of 200 genes was assigned a higher expression proportion of 90% in cluster 0, with a 70% expression proportion in the remaining clusters.

### Pathway enrichment analysis

Pathway enrichment analysis of DEGs was performed using the R package clusterProfiler(25) v4.7.1.003, covering both Gene Ontology (GO) and Kyoto Encyclopedia of Genes and Genomes (KEGG) analysis. Pathways with a *P* value less than 0.05 were considered significantly enriched. The GO categories, including molecular function (MF), biological process (BP), and cellular component (CC) categories, were used as references.

### Differentially accessible regions enrichment using SNPsea

We used single nucleotide polymorphism set enrichment analysis (SNPsea(26)) to analyze the enrichment of differentially accessible regions in different tissues and cell types. The SNPs corresponding to the differentially accessible regions were mapped to the reference genome, followed by enrichment analysis using SNPsea to determine whether they are significantly concentrated in specific tissues or cell types. Next, we used SNPsea calculated enrichment scores and assessed statistical significance by comparing the input SNP set with known SNP sets associated with tissue- or cell type-specific gene expression. The final results revealed the tissues or cell types that these differentially accessible regions may be involved in regulating, providing clues for further understanding their function in specific biological contexts.

### Peak annotation and enrichment analysis using ChIPseeker

We used the R package ChIPseeker(27) v1.34.1 to annotate differentially accessible regions within functional regions of the genome and analyze their enrichment. First, the *annotatePeak* function was employed to annotate the differentially accessible regions in the genome, determining their positions within the genome, such as promoter regions, introns, exons, or downstream regions. Next, the *plotAnnoBar* function was used to visualize the genomic distribution of the differentially accessible regions, providing an intuitive display of region enrichment across different functional regions.

### Evaluation of computational efficiency of CSFeatures

We utilized simulated scRNA-seq data generated by splatter v1.28.0, including five cell populations, to measure time and memory usage for running CSFeatures. Experiments were conducted on an Intel(R) Xeon(R) Platinum 8375C CPU. The complexity of our method is closely related to the number of cells and genes. To assess the impact of gene quantity, we fixed the cell count at 10,000 and varied the gene count from 4,000 to 20,000 in increments of 4,000. Similarly, to evaluate the effect of cell quantity, we fixed the gene count at 20,000 and varied the cell count from 10,000 to 50,000 in increments of 10,000.

### Statistical analysis

Unless otherwise specified, the Wilcoxon rank-sum test is used to identify differentially expressed genes for scRNA-seq and ST data, while LR is used to identify differentially accessible regions for scATAC-seq and spaATAC-seq data. Results are ranked by *P* values, adjusted for multiple testing correction using the Benjamini-Hochberg, with differences considered statistically significant at *P* < 0.01.

## Results

### Limitations of existing differential analysis methods in single-cell and spatial data

In scRNA-seq data analysis, differential gene expression analysis is essential for tasks such as cell type annotation, functional enrichment of cell populations, and investigating functional changes under different conditions(28) (Figure 1A). However, we identified several common shortages in current differential analysis (DA) methods for recognizing differentially expressed genes (DEGs). First, in conducting differential analysis, the target cell population is compared against all other cells combined. This approach can lead to situations where a gene may be highly expressed in a few cell populations within the “other” group but has low overall expression across the entire “other” group, resulting in a low *P* value and a high ranking. The Wilcoxon rank-sum test is one of the most commonly used methods and is also the default in popular analysis frameworks such as Seurat(9) and Scanpy(10). Given this, we used the Wilcoxon rank-sum test to identify DEGs in the scRNA-seq dataset from tumor and adjacent non-tumor tissues of gastric cancer patients (referred to as the gastric cancer dataset)(29). For example, we selected fibroblasts as the target cell population, we found that among the top 20 ranked genes, only one gene, namely *MFAP4*, demonstrated clear cell type specificity (Supplementary Figure S1A). In contrast, other top-ranked genes, such as *DCN* (ranked first), were not only highly expressed in fibroblasts but also exhibited substantial expression in mural cells (Figure 1B, C). However, despite being one of the most cell type-specific genes, *MFAP4* was ranked only eighth (Figure 1A). Second, some genes indeed show higher expression in the target cell population compared to the “other” group, leading to low *P* values and high rankings. However, these genes also exhibit considerable expression across all other cell populations, which undermines their cell type specificity. This issue was further validated using the peripheral blood mononuclear cells (PBMC) dataset profiled by 10x Genomics Chromium(9). For instance, in the DA results for B cells, we observed that the significantly differentially expressed gene *CD74* was highly expressed not only in the target population but also across all other cell populations (Figure 1D, E).

**Figure 1.**
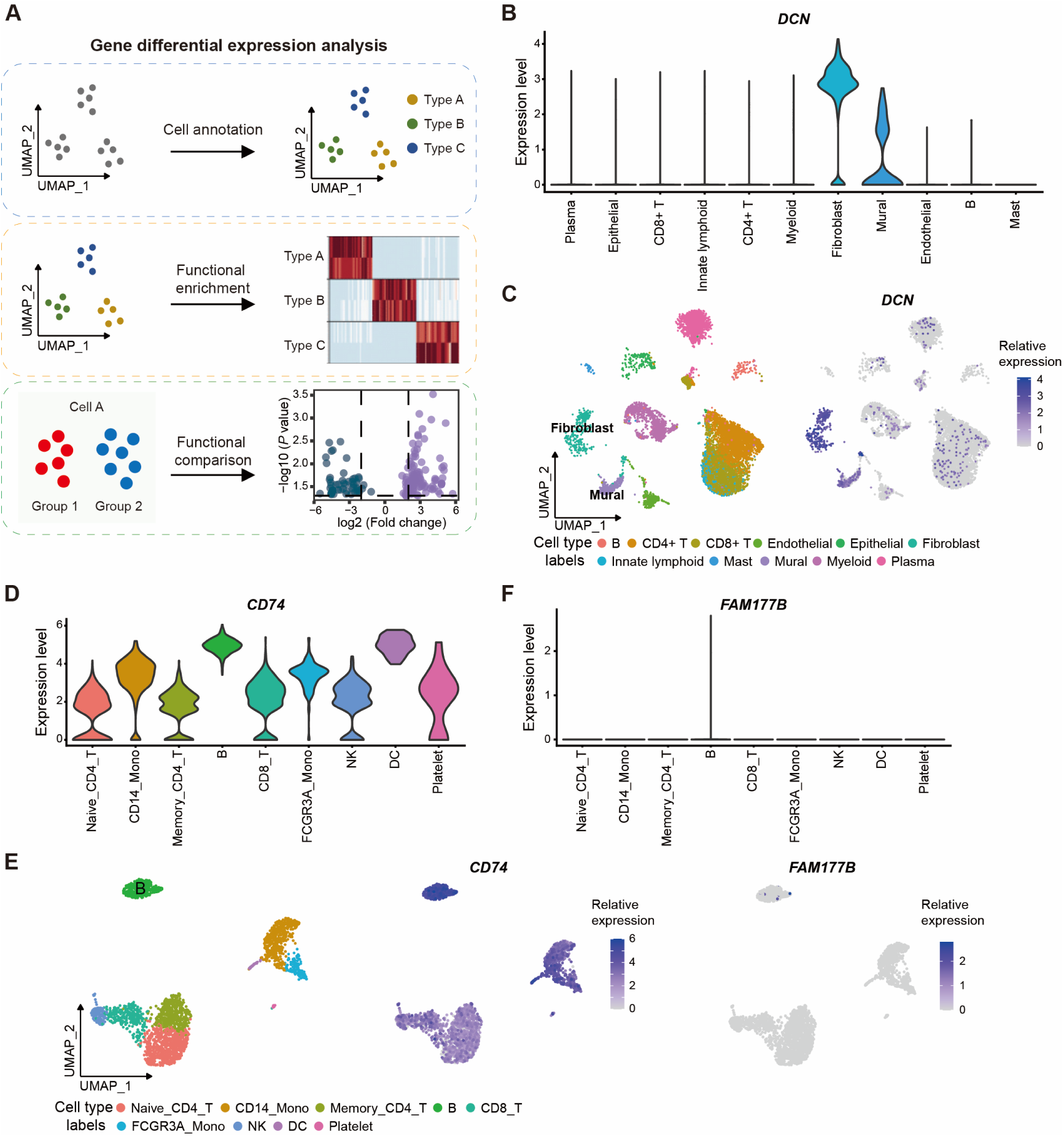
Limitations of differential expression analysis in scRNA-seq data. (**A**) Application scenarios of differential expression analysis in scRNA-seq data analysis. (**B**) The violin plot shows the expression distribution of the top-ranked differentially expressed gene (DEG), *DCN*, identified by the Wilcoxon rank-sum test for the fibroblast population in the gastric cancer dataset. While highly expressed in fibroblasts, *DCN* also exhibits significant expression in mural cells. (**C**) The scatter plots visualize the expression distribution of the gene *DCN* across different cell populations within the gastric cancer dataset. (**D**) The violin plot illustrates the expression distribution of the gene *CD74*, identified by the Wilcoxon rank-sum test, for the B cell population in the PBMC dataset. *CD74* is highly expressed in B cells as well as in several other cell populations. (**E**) The scatter plots demonstrate the distribution of *CD74* and *FAM177B* gene expression across different cell populations in the PBMC dataset. (**F**) The violin plot illustrates the expression distribution of the top-ranked DEG, *FAM177B*, for the B cell population in the PBMC dataset. Identified through fold change ranking after filtering with *P* < 0.01, *FAM177B* exhibits low expression across all cell populations, including the B cell population.

Moreover, similar challenges are encountered in the analysis of scATAC-seq, ST, and spaATAC-seq data. For instance, in the scATAC-seq data from mixed human cell lines(30), only three of the top 20 differentially accessible regions identified by the LR method demonstrated cell type specificity in the K562 cell population (Supplementary Figure S2A). Similarly, the limitation of commonly used methods in lacking domain specificity when identifying differential features is evident in the Cortex_1 population of the mouse brain ST data (Supplementary Figure S3A) and the Cartilage_2 population from the mouse embryos spaATAC-seq data(31) (Supplementary Figure S3B).

Furthermore, another important metric in differential analysis is the fold change (FC). Researchers may set a threshold based on the *P* value from statistical tests (e.g., *P* < 0.01) to filter for significant DEGs, and then rank these genes based on their FC, focusing on those with high FC values. However, our investigation revealed that some genes identified using this approach, while showing very low expression in other cell populations, may also exhibit only weak expression in the target cell population. For example, in the DA results for B cells in the PBMC dataset, when DEGs with *P* < 0.01 were ranked in descending order of FC, the top-ranked gene, *FAM177B*, showed zero expression in all other cell populations. However, it was expressed in only 4.87% of the target B cell population (Figure 1E, F). Additionally, we show that the top 12 genes identified using this approach all exhibited poor performance, with only a small fraction of cells within the B cell population expressing these genes (Supplementary Figure S4). This outcome may be attributed to the fact that the FC-based strategy overlooks a critical factor: the proportion of gene expression within cell populations. Although some genes may exhibit higher expression intensity in the target cell population compared to other populations, their actual expression proportion may remain low. This omission can lead to incomplete or inaccurate identification of DEGs, as the expression intensity alone does not fully capture the biological significance of expression patterns across cell populations.

In conclusion, the top-ranked features identified by commonly used methods often lack cell type specificity. This issue is not only observed in scRNA-seq data but also in scATAC-seq and spatial omics data, regardless of whether the features are ranked by *P* value or fold change.

### CSFeatures identifies features with cell type specificity in differential analysis

To address the aforementioned limitations, we propose CSFeatures, a method for identifying cell type-specific differential features. Taking scRNA-seq data as an example, CSFeatures takes a gene expression matrix and cell population labels as input. First, the high-dimensional gene expression matrix is reduced to a low-dimensional space through a flexible dimensionality reduction method, with principal component analysis (PCA) as the default. Subsequently, a correlation matrix between cells is calculated using the low-dimensional representation, with Euclidean distance as the default, while allowing flexibility to incorporate other correlation measures as needed (Figure 2A and Methods). To avoid the issues of highly imbalanced sample sizes and increased biological heterogeneity within the “other” group, for each gene, CSFeatures comprehensively considers its average expression level, the smoothness of its expression distribution, and the proportion of cells expressing the gene within each cell population. We devised the expression index (EI) metric to quantify the degree of cell type specificity for each gene. Specifically, the EI value integrates information from five aspects simultaneously: the average expression of the gene in the target cell population (referred to as *M_t_*), the smoothness of its expression distribution in the target cell population (referred to as *S_t_*), the proportion of cells expressing the gene in the target cell population (referred to as *P_t_*), the local maximum expression in the “other” cell populations (referred to as *L_o_*), and the proportion of cells expressing the gene in the “other” cell populations (referred to as *P_o_*) (Figure 2A and Methods). Among these, *L_o_* is defined as the average expression level of the gene in the top *N* neighboring cells within the “other” populations. *P_o_* is calculated by aggregating the expression proportions across each cell population within the “other” group (Methods). Genes with higher *M_t_*, more uniform *S_t_*, and higher *P_t_* in the target cell population, combined with lower *L_o_* and *P_o_* in the “other” populations, will have higher EI values. Finally, genes are ranked by their EI values, prioritizing those with greater cell type specificity (Figure 2A and Methods).

**Figure 2.**
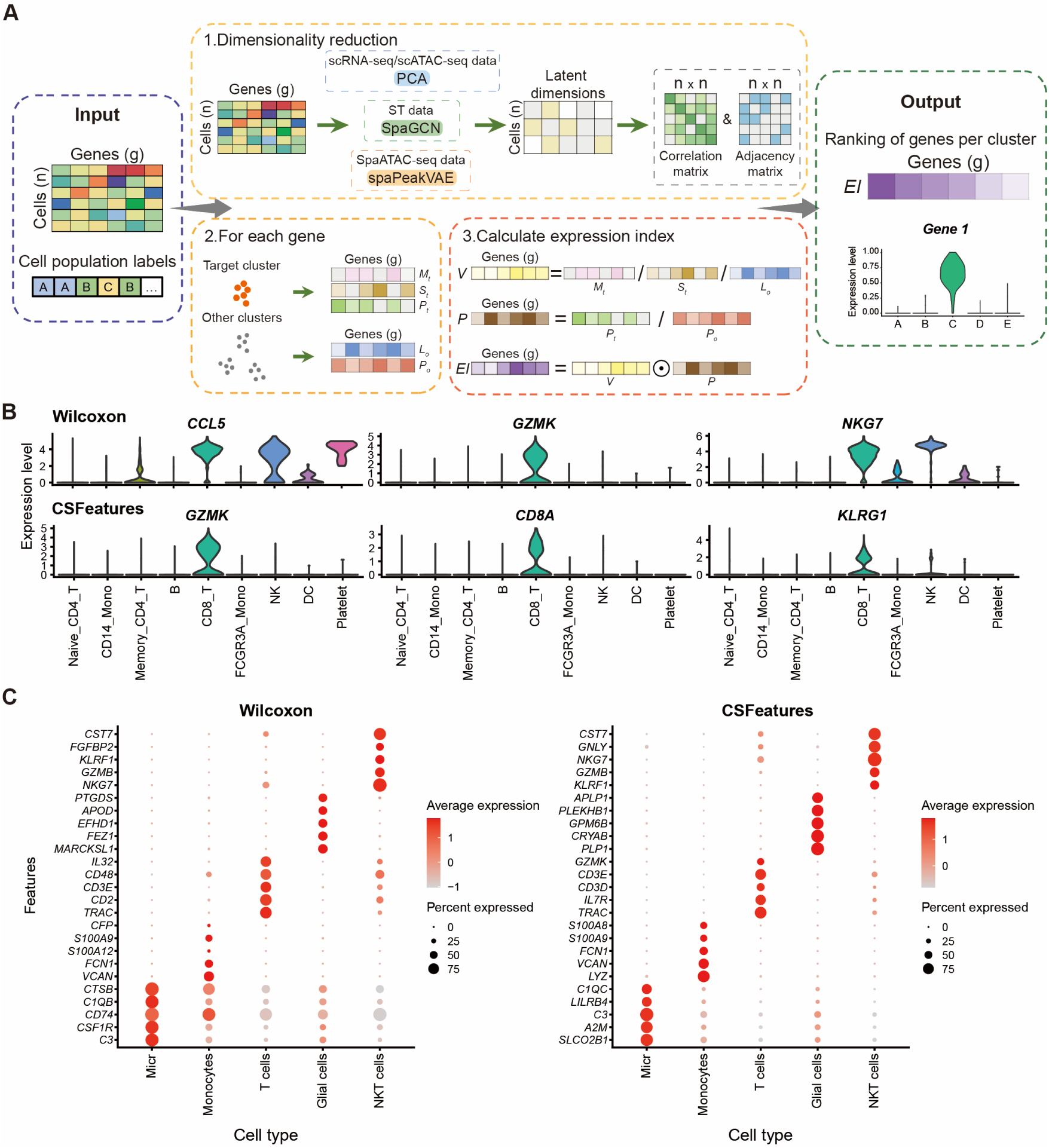
CSFeatures identifies cell type-specific differentially expressed genes in scRNA-seq data. (**A**) Overview of CSFeatures. CSFeatures takes a gene expression matrix and cell population labels as input, and computes a cell-to-cell correlation matrix following dimensionality reduction. For each gene, CSFeatures fully considers its expression level, the smoothness of its expression distribution, and the proportion of cells expressing the gene across all cell populations. Genes are ranked by their EI values, prioritizing those with strong cell type specificity. (**B**) For the CD8 T cell population in the PBMC dataset, the expression distribution of the top three genes identified by the Wilcoxon rank-sum test (top) and CSFeatures (bottom). (**C**) The bubble plots display the top five differentially expressed genes for each cell population identified by the Wilcoxon rank-sum test (left) and CSFeatures (right) in the human glioblastoma data. Colors represent expression levels, and bubble sizes correspond to the proportion of cells expressing each gene.

Moreover, CSFeatures is applicable to differential feature identification across multiple omics data, including scRNA-seq, scATAC-seq, ST, and spaATAC-seq data. For dimensionality reduction, CSFeatures allows flexible selection of methods. Specifically, PCA is used by default for preprocessed scRNA-seq and scATAC-seq data. In contrast, to fully incorporate spatial coordinates in spatial omics data, default methods such as SpaGCN(13) for ST data and spaPeakVAE(22) for spaATAC-seq data are integrated, though other methods are also supported (Figure 2A and Methods).

To demonstrate the effectiveness of CSFeatures, we first evaluated the performance in identifying cell type-specific DEGs from scRNA-seq data. Compared to the Wilcoxon rank-sum test, CSFeatures consistently assigned higher rankings to genes with greater cell type specificity. For example, for the fibroblasts in the gastric cancer dataset, only one of the top 20 DEGs identified by the Wilcoxon rank-sum test, *MFAP4*, showed high cell type specificity (Supplementary Figure S1A). In contrast, the majority of the top 20 genes identified by CSFeatures demonstrated significant specificity to fibroblasts (Supplementary Figure S1B). Moreover, for the myeloid cells in the gastric cancer dataset, half of the top 20 DEGs identified by the Wilcoxon rank-sum test performed poorly (Supplementary Figure S5A), whereas all of the top 20 DEGs identified by CSFeatures exhibited high cell type specificity (Supplementary Figure S5B). Notably, nine DEGs were identified by both methods, all of which demonstrated strong myeloid cell specificity. Beyond the solid tumor datasets, we observed consistent findings in the scRNA-seq datasets from leukemia patients(32) (Supplementary Figure S6) and mouse models of multiple myeloma(33) (Supplementary Figure S7). Compared to the Wilcoxon rank-sum test, CSFeatures proved more effective at identifying cell-type-specific differentially expressed genes. Furthermore, we validated this conclusion using the simulated scRNA-seq dataset (Supplementary Figure S8 and Methods).

Additionally, similar findings were observed for the CD8 T cells in the PBMC dataset. Among the top three genes identified by Wilcoxon rank-sum test, only *GZMK* exhibited CD8 T cell specificity, while the other two showed high expression across some of other cell populations (Figure 2B). In contrast, after ranking by CSFeatures, the top three genes, *GZMK*, *CD8A*, and *KLRG1*, all demonstrated strong cell type specificity (Figure 2B). Moreover, beyond the Wilcoxon rank-sum test, we evaluated other commonly used DA methods, including bimod(23), Student’s t-test, negbinom, Poisson, logistic regression (LR), and MAST(7). Taking the CD8 T cell population in the PBMC dataset as an example, we found that *CCL5* frequently ranked first across most of these methods, despite being highly expressed in multiple cell populations (Supplementary Figure S9). Notably, none of these methods were able to rank genes by cell type specificity as effectively as CSFeatures. Furthermore, we tested the effectiveness of CSFeatures on scRNA-seq data profiled by different sequencing protocols, including the human hepatocellular carcinoma dataset profiled by the 10x Genomics Chromium (referred to as the HCC dataset)(34), the mouse kidney dataset profiled by Drop-seq(35), and the human glioblastoma dataset profiled by the CEL-seq2(36). Consistent results across different datasets and cell populations demonstrate that CSFeatures more effectively identifies cell type-specific DEGs compared to commonly used DA methods (Figure 2C; Supplementary S10). In brief, CSFeatures effectively addresses the challenge of insufficient cell type specificity in DEG identification.

### CSFeatures facilitates more precise downstream analysis of scRNA-seq data

The identification of DEGs is of significant importance in scRNA-seq data analysis, as it provides key insights into the functional states that are unique to distinct cell populations(37,38). We have previously validated that the DEGs identified by CSFeatures exhibit cell type specificity in their expression patterns. To further explore whether these genes also reflect the functional characteristics of the target cell population, we first examined cell populations that have been extensively studied. Using the B cell population in the PBMC dataset as an example, we identified DEGs using both the Wilcoxon rank-sum test and CSFeatures. We compared the top 10 DEGs from each method to observe their correspondence and rank changes. The results showed that CSFeatures elevated the rankings of certain B cell type-specific DEGs, such as *TCL1A* and *VPREB3* (Figure 3A; Supplementary Figure S11). Furthermore, genes that already exhibited strong B cell type specificity, such as *CD79A* and *MS4A1*, remained highly ranked in the results generated by CSFeatures (Figure 3A; Supplementary Figure S11). Notably, the genes elevated in the ranking by CSFeatures have been reported to be closely associated with B cell function and activation(39,40). Additionally, in the analysis of fibroblasts within the gastric cancer dataset, we found that the top 10 genes uniquely identified by CSFeatures were all associated with fibroblast function (Supplementary Table S1). In contrast, some of the genes uniquely identified by the Wilcoxon rank-sum test, which did not exhibit the expected fibroblast-specific expression, had not been previously reported to have direct associations with fibroblasts either (Supplementary Figure S1A; Supplementary Table S2). These findings suggest that CSFeatures holds a distinct advantage in identifying genes related to the specific functions of particular cell populations. Given this, CSFeatures presents itself as a valuable tool for discovering marker genes that define distinct cell populations, providing valuable insights for studies focusing on cell identity and functional characterization.

**Figure 3.**
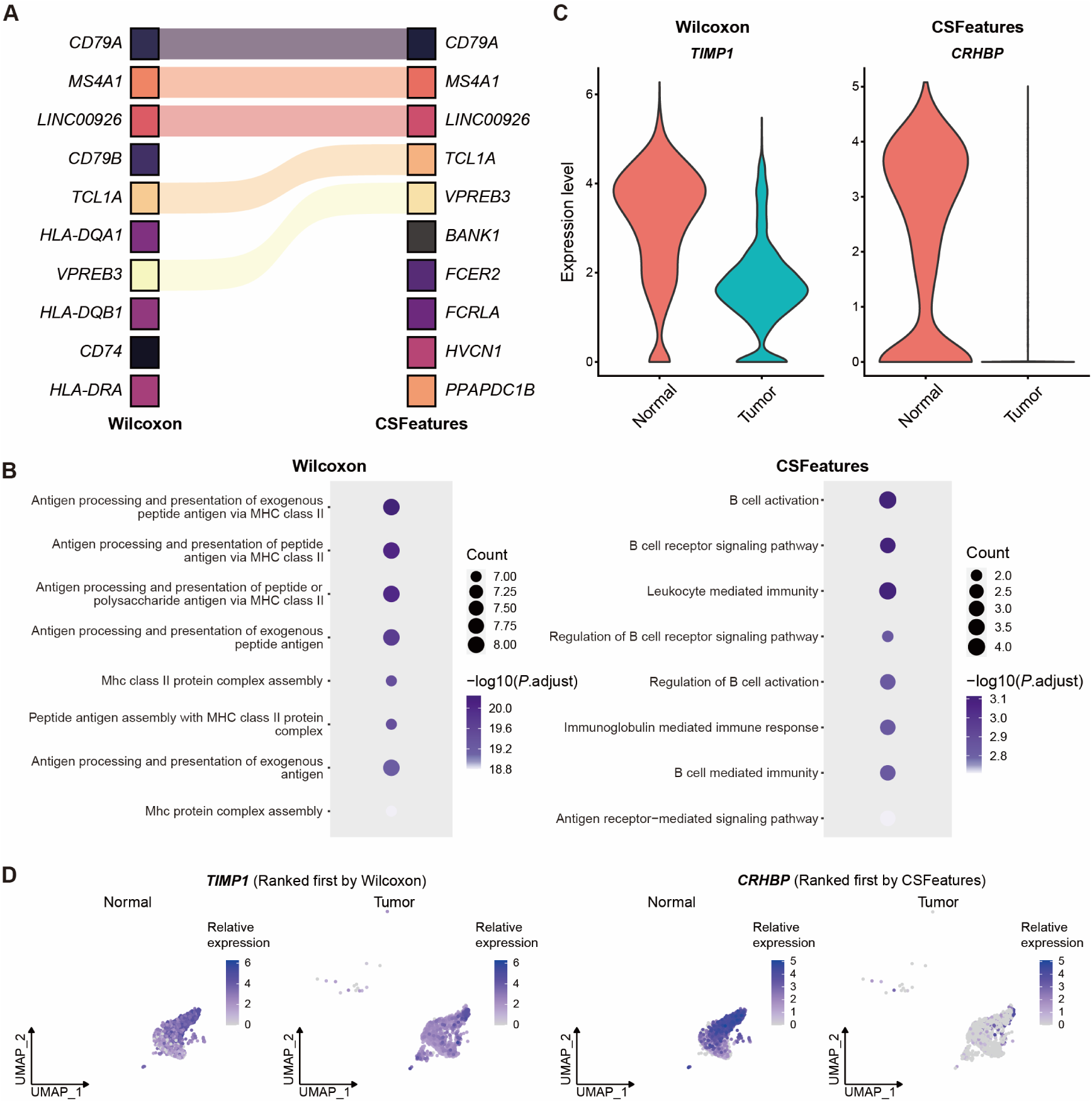
CSFeatures facilitates more precise downstream analysis of scRNA-seq data. (**A**) The top 10 differentially expressed genes (DEGs) identified by the Wilcoxon rank-sum test (left) and CSFeatures (right) for the B cell population in the PBMC dataset. (**B**) GO enrichment of the top 20 unique DEGs identified by the Wilcoxon rank-sum test (left) and CSFeatures (right) for the B cell population in the PBMC dataset. (**C**) For the endothelial cell population in the lung cancer dataset, the top downregulated genes in tumor tissue, *TIMP1* and *CRHBP*, respectively identified by the Wilcoxon rank-sum test (left) and CSFeatures (right), are shown. (**D**) The scatter plots further show the expression distribution of *TIMP1* and *CRHBP* between the groups, with color intensity representing expression levels.

To investigate the relationship between genes and their associated functions, we further conducted gene enrichment analysis. For the target cell population, we selected the top 20 genes uniquely identified by the Wilcoxon rank-sum test and CSFeatures, respectively. We then performed the Gene Ontology (GO) enrichment analysis on the unique genes identified by each method(25). We found that the cell type-specific DEGs identified by CSFeatures exhibited more precise functional annotations. Specifically, for B cells in the PBMC dataset, the top-ranked genes uniquely identified by the Wilcoxon rank-sum test were more likely to be enriched in broad terms, such as “antigen processing and presentation of exogenous peptide antigen via MHC class II” (Figure 3B). In contrast, the genes ranked by CSFeatures were more likely to be enriched in terms directly related to the specific functions of the target cell population, such as “B cell activation” and “B cell receptor signaling pathway” (Figure 3B). This further demonstrates that, compared to commonly used DA methods, CSFeatures is more effective in identifying DEGs associated with cell type-specific functional states.

Another key application of CSFeatures is identifying DEGs within a specific cell population under different conditions. For example, for the endothelial cells in the HCC dataset, we identified DEGs between tumor and adjacent non-tumor tissues using both the Wilcoxon rank-sum test and CSFeatures. Notably, among the top 10 genes identified as specifically expressed in the normal tissues by the Wilcoxon rank-sum test, several genes also showed high expression levels in tumor tissues (Supplementary Figure S12A). For clarity, in this context, DEGs in the target cell population from normal tissues are considered to be downregulated in tumor tissues compared to normal tissues. Conversely, DEGs in the target cell population from tumor tissues are considered to be upregulated in tumor tissues compared to normal tissues. This inconsistency highlights the issue that commonly used DA methods lack cell type specificity. Specifically, among the downregulated genes in tumor tissues, *TIMP1*, ranked first by the Wilcoxon rank-sum test, was downregulated compared to non-tumor tissues but still showed relatively broad expression in tumor tissues (Figure 3C, D). In contrast, *CRHBH*, ranked first by CSFeatures, exhibited extremely low expression levels in tumor tissues (Figure 3C, D). Moreover, this pattern was also confirmed in the analysis of the gastric cancer dataset. For example, in fibroblasts, among the top 10 genes identified by traditional methods as being upregulated in tumor tissues compared to normal tissues, genes such as *COL1A1* and *COL3A1* also exhibited high expression in the normal tissues. In contrast, the top 10 upregulated genes identified by CSFeatures were all specifically highly expressed in tumor tissues (Supplementary Figure S12B).

In summary, the genes identified by CSFeatures can clearly characterize the functional attributes of cell populations and reveal the heterogeneity among them. Furthermore, CSFeatures effectively identifies more specific DEGs between disease-associated cell states within the same population, offering valuable insights for understanding disease mechanisms.

### CSFeatures optimizes the identification of cell type-specific epigenetic features in scATAC-seq data

We next evaluated the performance of CSFeatures across other omics data. scATAC-seq provides valuable insights into gene regulation, chromatin states, and cell type-specific regulatory networks, and has been widely used in recent years(41–43). However, scATAC-seq data are typically high-dimensional and sparse(19,21). Currently, logistic regression (LR), which can detect significant differences even in low-signal backgrounds, has become a common method for identifying differentially accessible regions in scATAC-seq data(11). Nevertheless, regions identified by LR still suffer from the issue of lacking cell type specificity, similar to the challenges in differential analysis of scRNA-seq data. For example, in the mouse forebrain scATAC-seq data (referred to as the forebrain dataset)(44), we identified astrocyte (AC) cell type-specific regions via different methods. The results showed that half of the top 12 regions identified by LR performed poorly in terms of cell type specificity (Figure 4A). In contrast, after ranking by CSFeatures, all of the top 12 regions exhibited strong cell type specificity (Figure 4A). Additionally, similar findings were observed in the dataset derived from a mixture of human cell lines(30). For instance, for the K562 cell population, only 3 of the top 20 regions identified by LR demonstrated cell type specificity (Supplementary Figure S2A), whereas all of the top 20 regions ranked by CSFeatures showed strong specificity (Supplementary Figure S2B).

**Figure 4.**
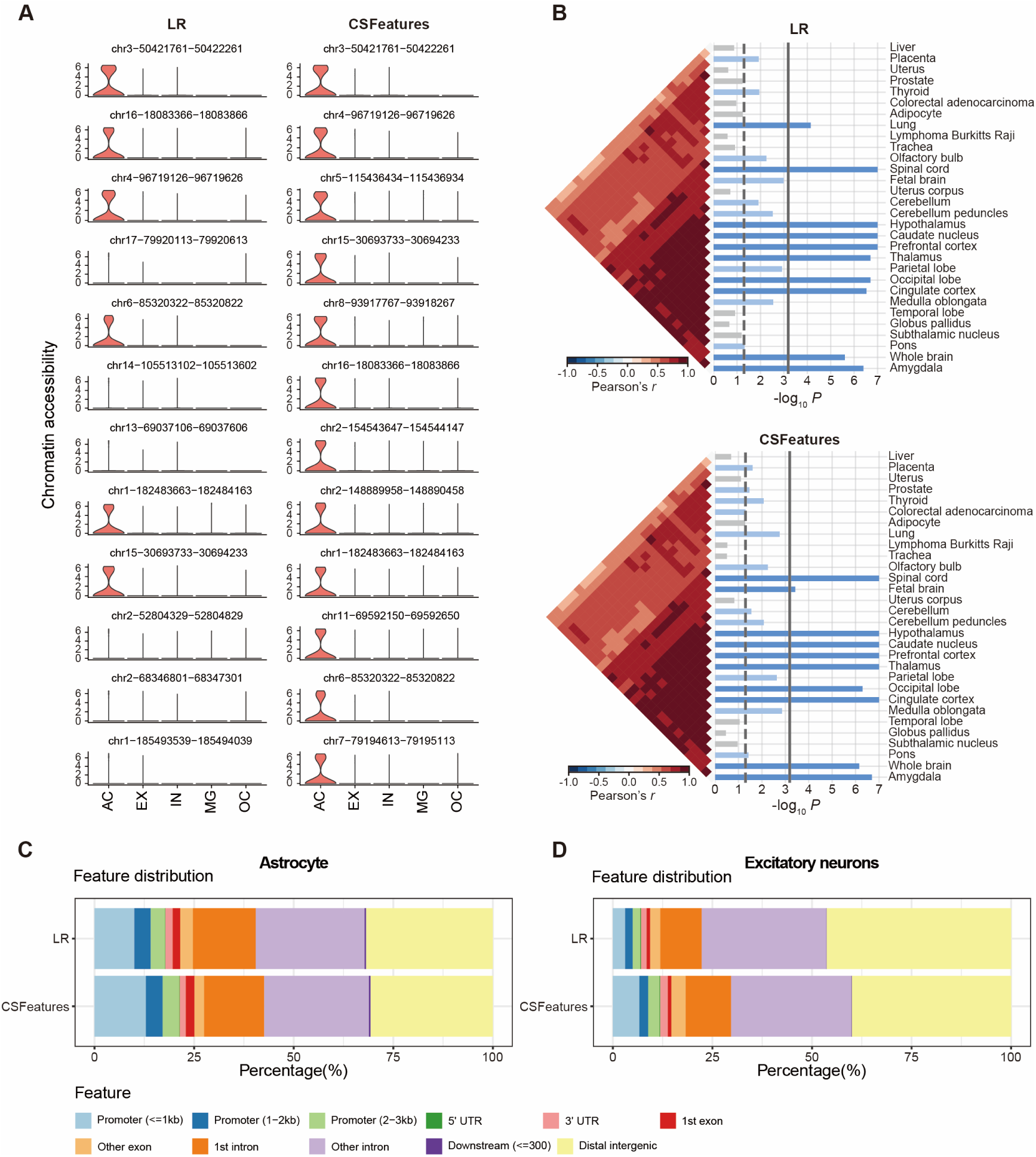
CSFeatures optimizes the identification of cell type-specific differentially accessible regions in scATAC-seq data. (**A**) The top 12 differentially accessible regions identified by LR (left) and CSFeatures (right) for the AC population in the forebrain dataset. (**B**) SNPsea analysis of the AC-specific differentially accessible regions identified by LR (top) and CSFeatures (bottom) shows the top 30 significantly enriched tissues. The vertical dashed and solid lines represent the unadjusted and Bonferroni-corrected one-sided *P* value cutoffs at 0.05, respectively. The heatmap illustrates the Pearson correlation coefficients of the expression profiles. (**C**) Utilizing ChIPseeker to visually present the distribution of astrocyte-specific differentially accessible regions identified by LR (top) and CSFeatures (bottom) across different genomic regions. (**D**) Utilizing ChIPseeker to visually present the distribution of excitatory neurons-specific differentially accessible regions identified by LR (top) and CSFeatures (bottom) across different genomic regions.

To further elucidate the biological significance of the differentially accessible regions, we selected the top 1,000 regions identified by LR and CSFeatures, respectively. We then conducted SNPsea analysis using default settings to examine their enrichment across different tissues(26). The results showed that regions identified by LR for AC cells derived from forebrain tissue were enriched in non-brain tissues, such as the lung tissue (Figure 4B). In contrast, regions identified by CSFeatures were exclusively enriched in brain-related tissues (Figure 4B). This highlights the potential of traditional methods to introduce confounding factors, whereas CSFeatures effectively mitigates this issue, identifying features with greater cell type specificity.

Furthermore, we examined the enrichment of these differentially accessible regions in functional genomic elements, which is a crucial step in understanding chromatin states, gene regulatory networks, and cellular functions(45). Differentially accessible regions enriched in promoter areas are typically associated with gene initiation and expression, while those enriched in enhancer areas may be involved in the regulation of gene expression. ChIPseeker can associate regions with genomic features such as promoters, exons, introns, and intergenic regions, generating detailed annotation results(27). We selected the top 1,000 differential regions identified by LR and CSFeatures, respectively, and used ChIPseeker to visually display their distribution across different genomic regions. Compared to the LR method, the differentially accessible regions selected by CSFeatures show a higher proportion of promoter enrichment (Figure 4C, D). This suggests that CSFeatures may more effectively filter out background noise, thereby more accurately capturing key gene regulatory regions.

Overall, CSFeatures has shown robust applicability in identifying cell type-specific differentially accessible regions in scATAC-seq data. CSFeatures facilitates the identification of key regulatory elements, uncovering critical biological mechanisms and offering valuable insights for epigenetic research.

### CSFeatures facilitates the identification of features with spatial patterns

In recent years, advancements in ST and spaATAC-seq technologies have allowed researchers to map gene expression and chromatin accessibility within tissue contexts, providing distinct yet complementary insights into spatial molecular regulation(31,46,47). The integration of spatial coordinates with omics data has proven essential for uncovering tissue heterogeneity and understanding the distinct roles of specific spatial domains(48,49). Key analytical steps in spatial omics include the identification of spatial domains, followed by the identification of differential features, such as gene expression or regulatory region accessibility, within these domains. However, while various methods exist to identify spatial features, they often lack specificity in capturing differential features tied to particular spatial domains, limiting their biological interpretability.

In ST data, DEGs between spatial domains can also identified by methods similar to those employed in scRNA-seq data analysis, such as the Wilcoxon rank-sum test or Student’s t-test. However, these approaches inherit the same limitations as in scRNA-seq data analysis, such as insufficient specificity in distinguishing features unique to particular spatial domains. Many methods developed for ST data are primarily designed to identify spatially variable genes (SVGs) by leveraging spatial coordinates and gene expression without the need for labeled information(13,15,16). While effective for detecting broad spatial patterns, these label-free approaches are not suited for identifying differentially expressed genes (DEGs) between distinct spatial domains. For example, when we tested SpaGCN(13) on the mouse brain ST data generated by the 10x Visium platform (referred to as the mouse brain dataset), its performance in identifying cell type-specific DEGs was also suboptimal. For the Thalamus_1 population, all of the top 12 genes identified by SpaGCN were not only highly expressed in Thalamus_1 but were also broadly expressed across other cell populations (Supplementary Figure S13).

While SpaGCN may fall short in achieving cell type specificity when identifying DEGs between spatial domains, it excels at integrating gene expression, spatial coordinates, and histological information using graph convolutional networks. CSFeatures utilizes the embeddings provided by SpaGCN to enable the effective identification of spatial domain-specific DEGs in ST data (Supplementary Text S1). We applied both the Wilcoxon rank-sum test and CSFeatures to the mouse brain dataset. Similar to SpaGCN, in the Thalamus_1 population, the Wilcoxon rank-sum test yielded suboptimal results. For example, the top-ranked genes, *Shox2* (ranked first) and *Synpo2* (ranked third), were highly expressed in both the target cell population and Thalamus_2 cells (Figure 5A, B). In contrast, the top three DEGs identified by CSFeatures were specifically and highly expressed in the Thalamus_1 population alone (Figure 5A, B).

**Figure 5.**
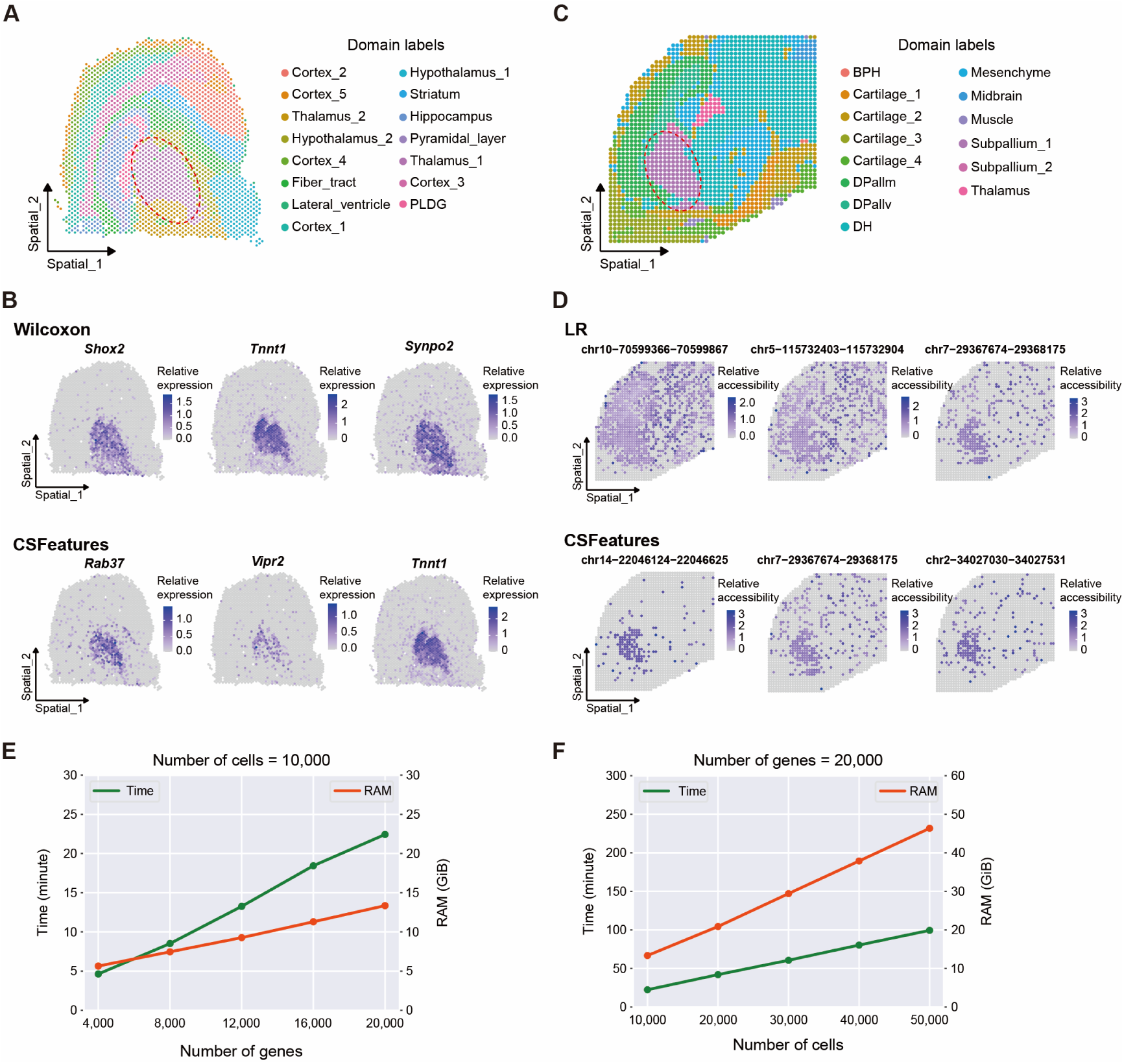
CSFeatures identifies features with distinct spatial profiles. **A**. Spatial domain annotation in the mouse brain ST dataset, with distinct colors representing different domains. **B**. Scatter plots illustrate the expression distribution of the top three differentially expressed genes identified by Wilcoxon rank-sum test (top) and CSFeatures (bottom) for the Thalamus_1 population in the mouse brain ST dataset. The color intensity reflects the gene expression levels. **C**. Spatial domain annotation in the mouse embryo spaATAC-seq dataset, with distinct colors representing different domains. **D**. Scatter plots illustrate the expression distribution of the top three differentially accessible regions identified by LR (top) and CSFeatures (bottom) for the Subpallium_1 population in the mouse embryo spaATAC-seq dataset. The color intensity reflects the chromatin accessibility levels. **E**. Using 10,000 cells as an example, the time and memory usage trends of running CSFeatures are shown as the number of genes increases. **F**. Using 20,000 genes as an example, the time and memory usage trends of running CSFeatures are shown as the number of cells increases.

Furthermore, CSFeatures is equally effective in identifying specific features in spaATAC-seq data. The spaATAC-seq technology enables the study of chromatin accessibility in situ within tissues, providing a novel epigenetic perspective for the emerging field of spatial biology(31). However, there are currently few tools specifically designed for spaATAC-seq data analysis. To fully integrate spatial coordinates when processing spaATAC-seq data, CSFeatures incorporates spaPeakVAE(22) in its dimensionality reduction step instead of PCA. Compared to the LR method, the differentially accessible regions identified by CSFeatures demonstrate superior domain specificity. For instance, in our analysis of the mouse embryo spaATAC-seq dataset(31), using Subpallium_1 population as an example, we observed that the top three differentially accessible regions identified by LR exhibited broad chromatin accessibility not only within the Subpallium_1 population but also across other cell populations within distinct spatial domains (Figure 5C, D). In contrast, the top three regions identified by CSFeatures demonstrated strong specificity for the Subpallium_1 population (Figure 5C, D). Furthermore, when examining the top 10 differentially accessible regions, we found that over half of those ranked by LR showed broad accessibility across most cell populations (Supplementary Figure S14A), whereas the regions identified by CSFeatures were predominantly specific to the Subpallium_1 population (Supplementary Figure S14B).

Overall, CSFeatures effectively integrates dimensionality reduction methods tailored for spatial omics data, enabling the integration of spatial coordinates while identifying cell type-specific features in both ST and spaATAC-seq data. This enhances our ability to interpret cellular heterogeneity and spatial gene regulation within tissues, offering valuable perspectives for understanding complex biological processes.

### Evaluation of the computational efficiency of CSFeatures

To assess the computational efficiency of our method, we utilized simulated scRNA-seq data generated by splatter(24) to measure the time and memory usage for running CSFeatures. We assessed resource consumption by fixing the cell count at 10,000 and increasing the gene count from 4,000 to 20,000 in increments of 4,000. Similarly, we evaluated the effect of varying cell counts by fixing the gene count at 20,000 and adjusting the cell count from 10,000 to 50,000 in increments of 10,000. The results demonstrate that our method maintains relatively low resource demands. Memory usage stays under 15 GB with 10,000 cells as the gene count reaches 20,000 (Figure 5E). Even with 50,000 cells and 20,000 genes, memory usage remains below 50 GB (Figure 5F). Furthermore, both memory and time consumption of CSFeatures scales almost linearly with both cell and gene counts (Figure 5E, F), ensuring its ability to handle large-scale computational tasks efficiently.

## Discussion

Advances in single-cell and spatial omics technologies now enable the precise analysis of gene expression, chromatin accessibility, and other molecular features at single-cell resolution, providing powerful tools to explore the complexity and heterogeneity of biological systems. A critical component of these analyses is the identification of differentially expressed genes or differentially accessible regions between cell populations or states, offering valuable insights into uncovering cellular functions and disease processes. This highlights the need for robust methods that can reliably detect significant variations between cell populations, driving further advancements in single-cell and spatial omics research.

Current differential analysis methods often fail to achieve the expected cell type specificity when identifying differential features for the target cell population. This is primarily because they fail to fully account for the strong heterogeneity present in other cell populations, leading to incomplete consideration of feature activity across all cell populations. Given the importance of identifying differential features and the limitations of current methods, we propose CSFeatures, a method designed to identify cell type-specific differential features. Taking scRNA-seq data as an example, for each gene, CSFeatures fully considers the gene expression levels, expression smoothness, and expression proportions in the target cell population as well as in each of the remaining cell populations. CSFeatures prioritizes genes that are highly expressed in the target population and lowly expressed elsewhere. We have extensively validated this conclusion using eight scRNA-seq datasets, including seven public datasets that were generated by different sequencing technologies from various species and different tissues, as well as one simulated dataset. Moreover, through extensive testing, we found that the underlying principle of CSFeatures can be generalized to other omics data, including scATAC-seq, ST, and spaATAC-seq data. Our results consistently show that CSFeatures effectively identifies cell type-specific molecular features, which exhibit functional enrichment closely aligned with the functional profiles of the corresponding target cell population. This capability enhances the understanding of cellular heterogeneity and contributes to uncovering underlying biological mechanisms.

Naturally, there remains potential for further optimization of CSFeatures. For instance, the computational demands increase with larger datasets containing numerous cells and genes. To address this, processing speed and memory utilization can be further optimized through the high-performance parallel computing frameworks. Moreover, CSFeatures currently identifies differential features based on provided cell clustering results, but we have found that certain cell populations inherently have relatively few cell type-specific features. We wonder whether cells that specifically express the same set of features should be grouped together. In the future, optimizing clustering results based on cell type-specific features could be an interesting direction to explore.

## Data availability

The PBMC scRNA-seq dataset can be downloaded at https://cf.10xgenomics.com/ samples/cell/pbmc3k. The human hepatocellular carcinoma scRNA-seq dataset can be retrieved from NCBI Gene Expression Omnibus (GEO) with accession number GSE149614. The human gastric cancer scRNA-seq dataset can be retrieved from GEO with accession number GSE206785. The human leukemia scRNA-seq dataset can be retrieved from GEO with accession number GSE207356. The mouse multiple myeloma scRNA-seq dataset can be retrieved from GEO with accession number GSE205393. The mouse kidney scRNA-seq dataset by Drop-seq can be retrieved from GEO with accession number GSE220045. The human glioblastoma scRNA-seq dataset by CEL-seq2 can be retrieved from GEO with accession number GSE166418. The mouse forebrain scATAC-seq dataset can be retrieved from GEO with accession number GSE100033. The mixture of human cell lines scATAC-seq dataset can be retrieved from NCBI Sequence Read Archive (SRA) with accession number SRP167062. The mouse brain ST dataset can be downloaded at https://www.10xgenomics.com/datasets/mouse-brain-section-coronal-1-standard-1-1-0. The mouse embryo spaATAC-seq dataset can be downloaded at https://www.10xgenomics.com/resources/datasets/flash-frozen-cortex-hippocampus-and-ventricular-zone-from-embryonic-mouse-brain-e-18-1-standard-1-2-0. Details of these datasets are described in Supplementary Table S3.

CSFeatures is fully interoperable with established single-cell and spatial omics data analysis workflows and is implemented as an open-source Python package, with documentation and tutorials available at https://github.com/xuyungang/CSFeatures.

## Supplementary data

Supplementary Data are available at NAR Online.

## Author contributions

R.L., Y.X. and S.C. were responsible for the design, analysis, organization, as well as drafting and revising the manuscript. S.C., R.L. and Y.L. designed, implemented, and validated CSFeatures. R.L., Y.L., H.H., Z.L., Y.Z., and Y.Y. participated in data collection and analysis. Y.X. and S.C. supervised the study.

