## Supplemental Data 1 for "CSFeatures improves identification of cell type-specific differential features in single-cell and spatial omics data"

### Supplementary Texts

#### Text S1: Selection of dimensionality reduction methods for CSFeatures in spatial transcriptomics data processing

In processing spatial transcriptomics (ST) data, CSFeatures integrates established dimensionality reduction methods to fully incorporate spatial coordinates. We evaluated the performance of CSFeatures after integrating STAGATE(1), SpatialPCA(2), and SpaGCN(3) respectively. Using the Thalamus_1 population in the mouse brain dataset as an example, all approaches demonstrated satisfactory performance overall. All methods consistently identified the top three genes as *Rab37*, *Tnnt1*, and *Vipr2*, with *Vipr2* demonstrating greater cell type specificity than *Tnnt1* (Fig 5d). Notably, the gene *Vipr2* ranked second in SpaGCN, while it ranked third in both STAGATE and SpatialPCA. Additionally, SpaGCN excels at integrating gene expression, spatial coordinates, and histological information through graph convolutional networks. Based on these findings, we chose SpaGCN as the default dimensionality reduction method for ST data.

### Supplementary Figures

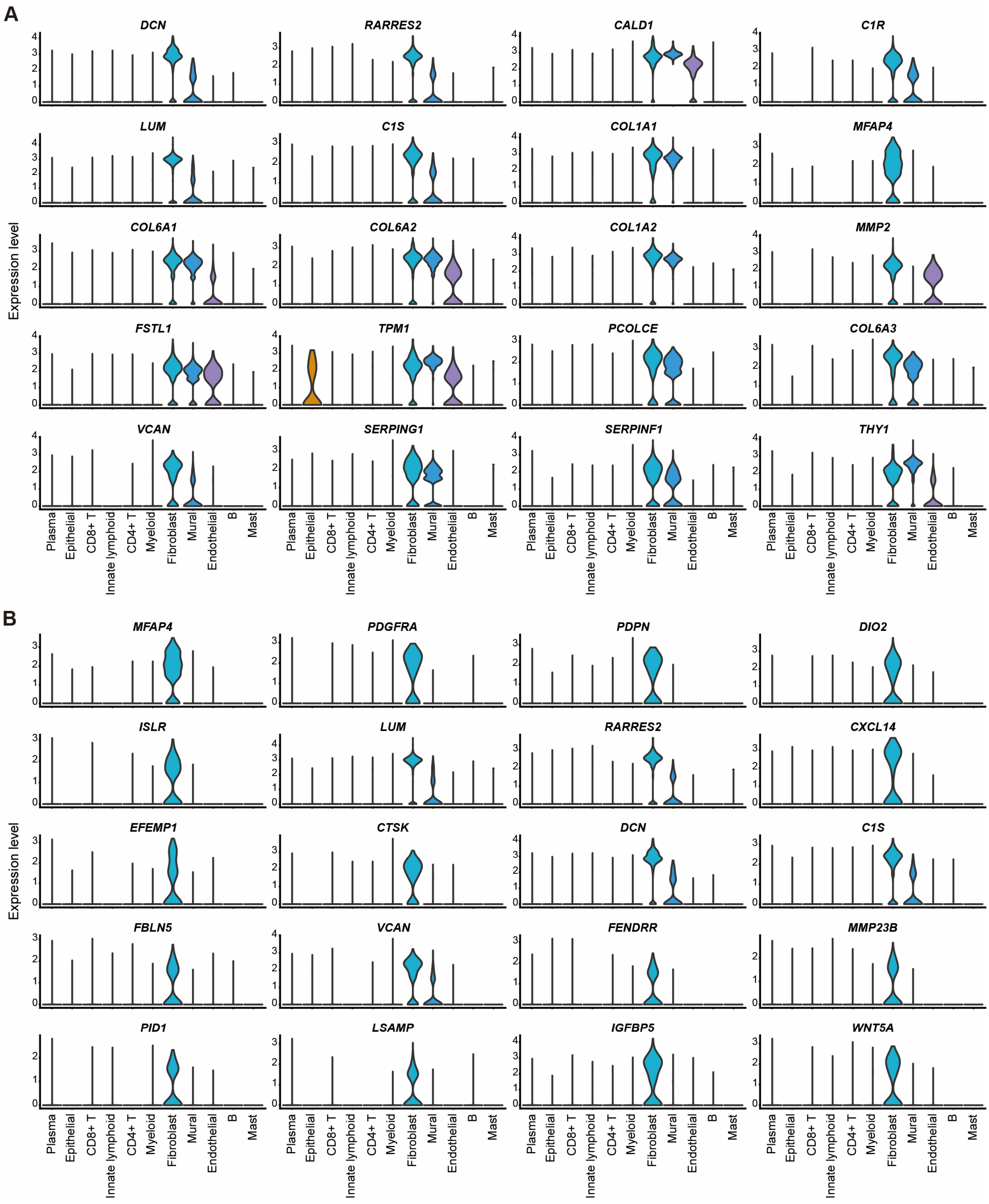

**Figure S1. The expression distribution of differentially expressed genes for the fibroblast population in the gastric cancer dataset.** (**A**) The violin plots show the expression distribution of the top 20 differentially expressed genes identified by the Wilcoxon rank-sum test across different cell populations. Among these, only the eighth-ranked gene, *MFAP4*, exhibits fibroblast-specific expression. (**B**) The violin plots show the expression distribution of the top 20 differentially expressed genes identified by CSFeatures across different cell populations. Fifteen of these genes exhibit strong fibroblast-specific expression.

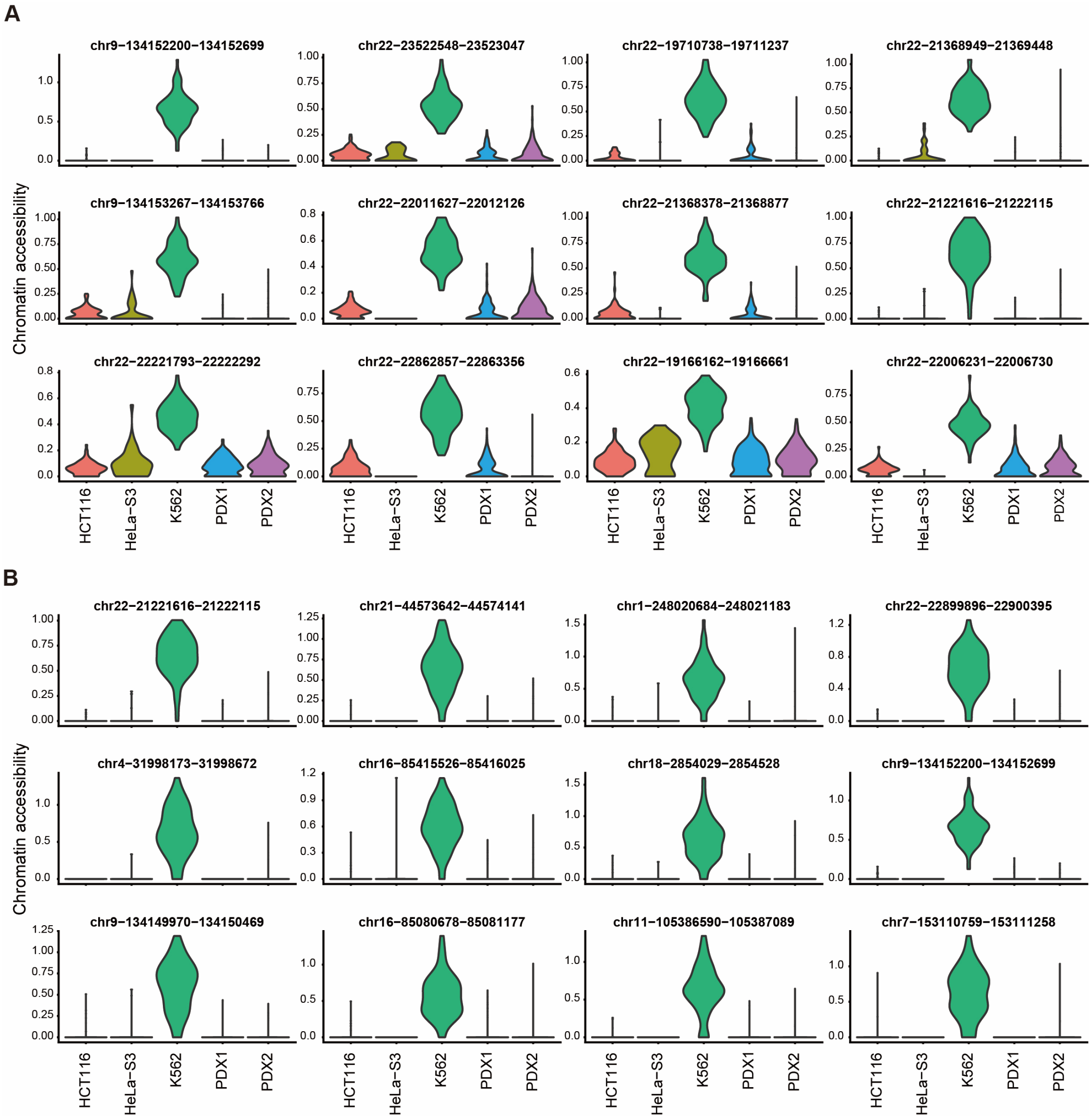

**Figure S2. Distribution of differentially accessible regions for the K562 cell population in the human cell line dataset.** (**A**) The distribution of the top 20 differentially accessible regions identified by LR across different cell populations, with only two regions exhibiting K562 cell type specificity. (**B**) The distribution of the top 20 differentially accessible regions identified by LR across different cell populations, with all regions exhibiting strong K562 cell type specificity.

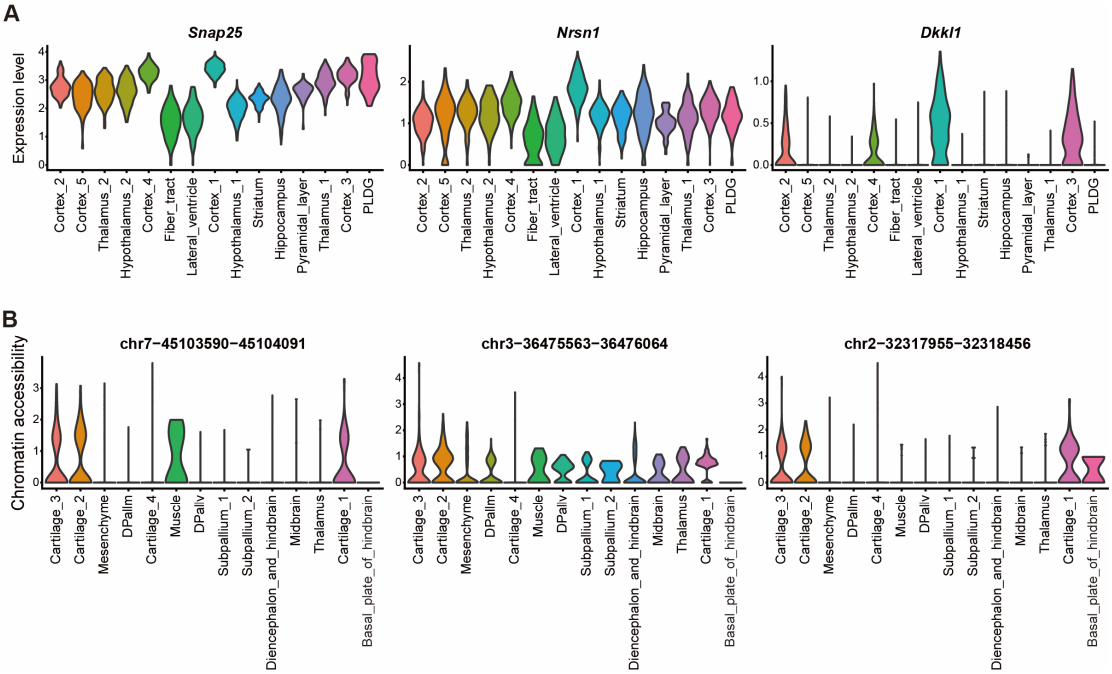

**Figure S3. The limitations of differential analysis in spatial omics data.** (**A**) The top three differentially expressed genes identified by the Wilcoxon rank-sum test for the Cortex_1 population in the mouse brain ST data exhibit poor cell type specificity. (**B**) The top three differentially accessible regions identified by LR for the Cartilage_2 population in the mouse embryo spaATAC-seq data exhibit poor cell type specificity.

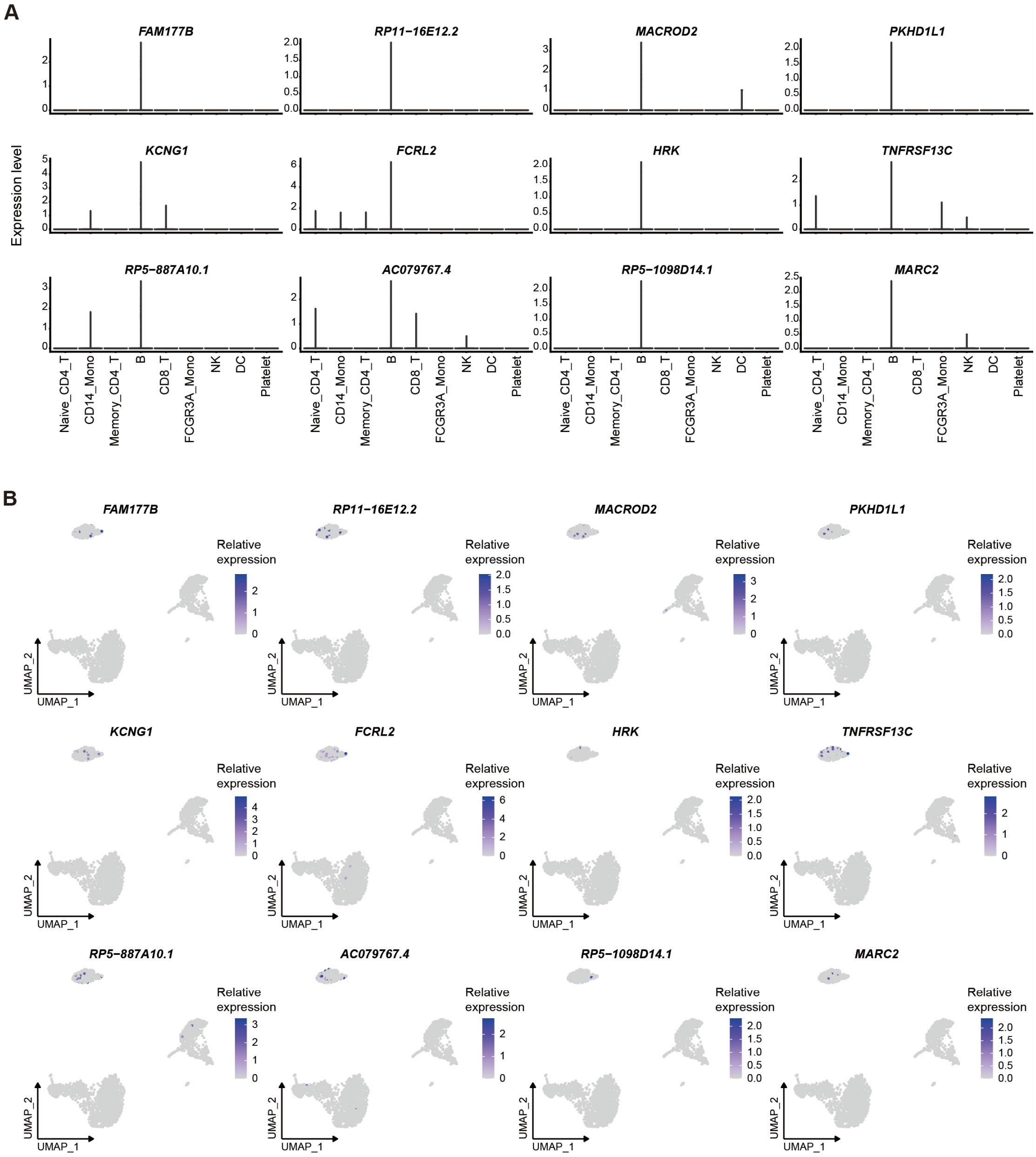

**Figure S4. The expression distribution of differentially expressed genes ranked by fold change for the B cell population in the PBMC dataset.** (**A**) The violin plots display the expression distribution of the top 12 differentially expressed genes, ranked by descending fold change after filtering with $P < 0.01$. All of these genes show only minimal expression in the target B cells. (**B**) The scatter plots further illustrate the expression distribution of the top 12 differentially expressed genes, ranked by descending fold change after filtering with $P < 0.01$. The color intensity reflects the gene expression levels.

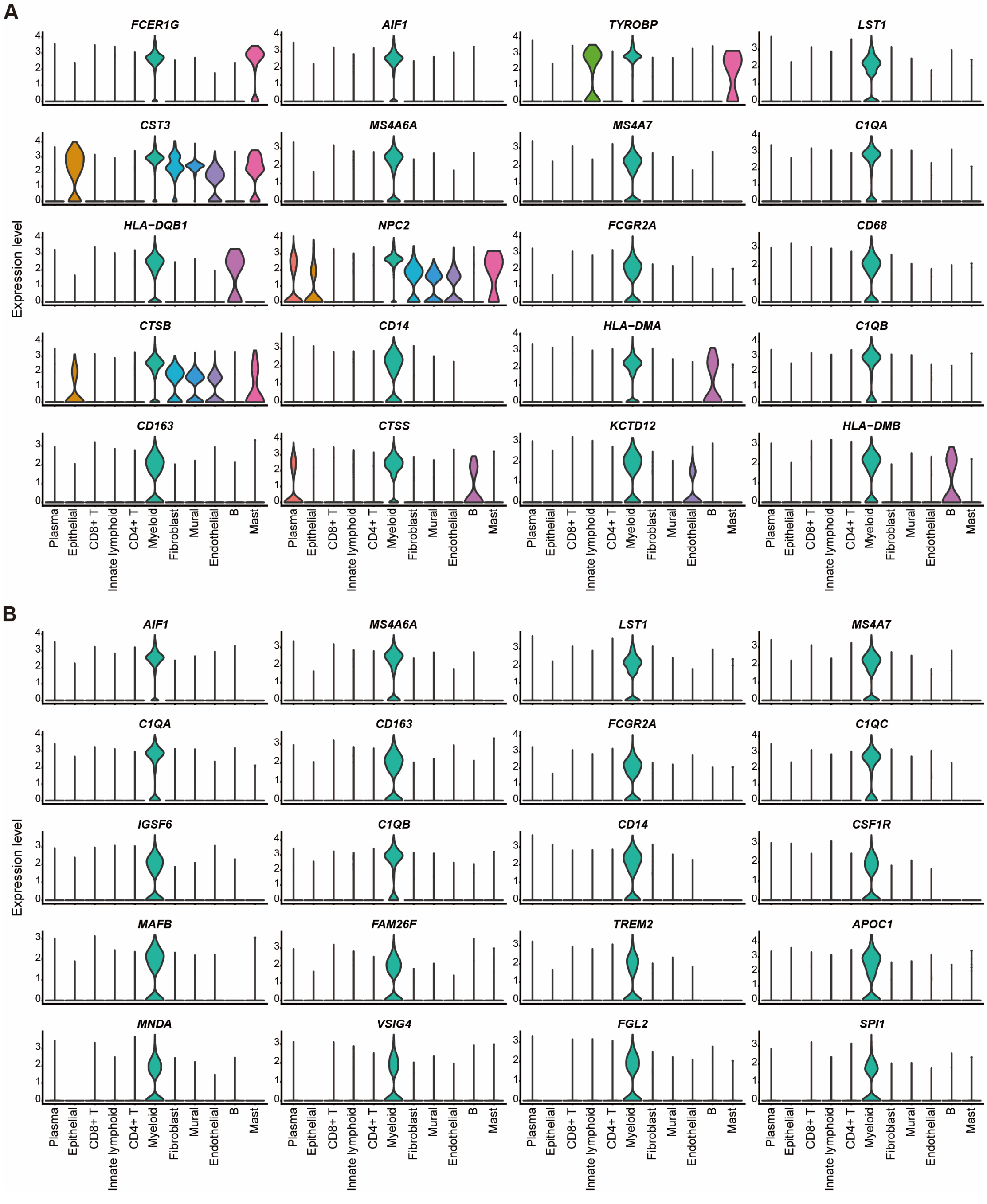

**Figure S5. The expression distribution of differentially expressed genes for the myeloid cell population in the gastric cancer dataset.** (**A**) The violin plots show the expression distribution of the top 20 differentially expressed genes identified by the Wilcoxon rank-sum test across different cell populations. Half of these genes lack specificity for myeloid cells. (**B**) The violin plots show the expression distribution of the top 20 differentially expressed genes identified by CSFeatures across different cell populations. All genes exhibit strong myeloid cell type-specific expression.

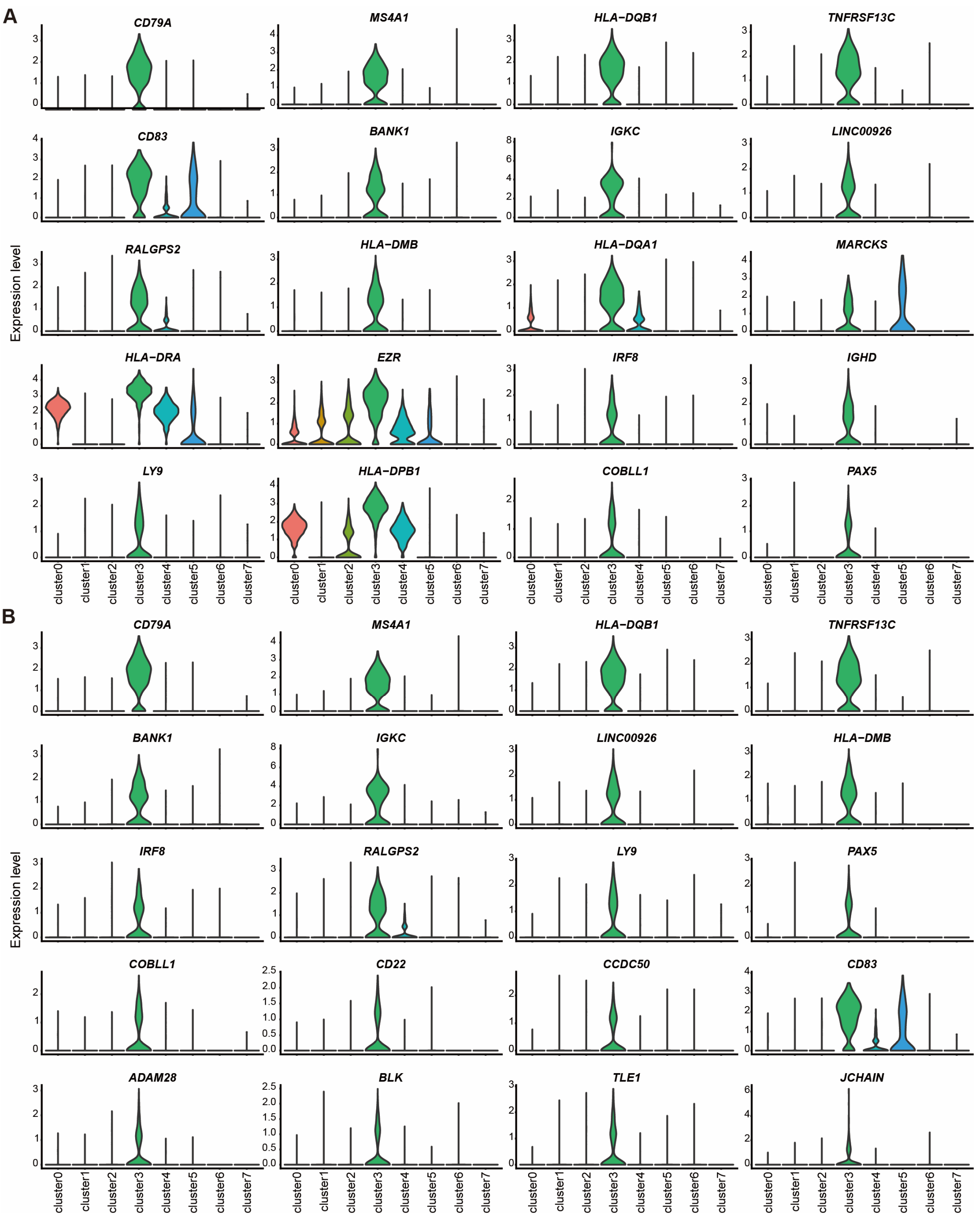

**Figure S6. The expression distribution of differentially expressed genes for the cluster 3 cell population in the scRNA-seq dataset from leukemia patients.** (**A**) The violin plots show the expression distribution of the top 20 differentially expressed genes identified by the Wilcoxon rank-sum test across different cell populations. Seven of these genes lack specificity for cluster 3 cell population. (**B**) The violin plots show the expression distribution of the top 20 differentially expressed genes identified by CSFeatures across different cell populations. Eighteen of these genes exhibit strong cell type-specific expression.

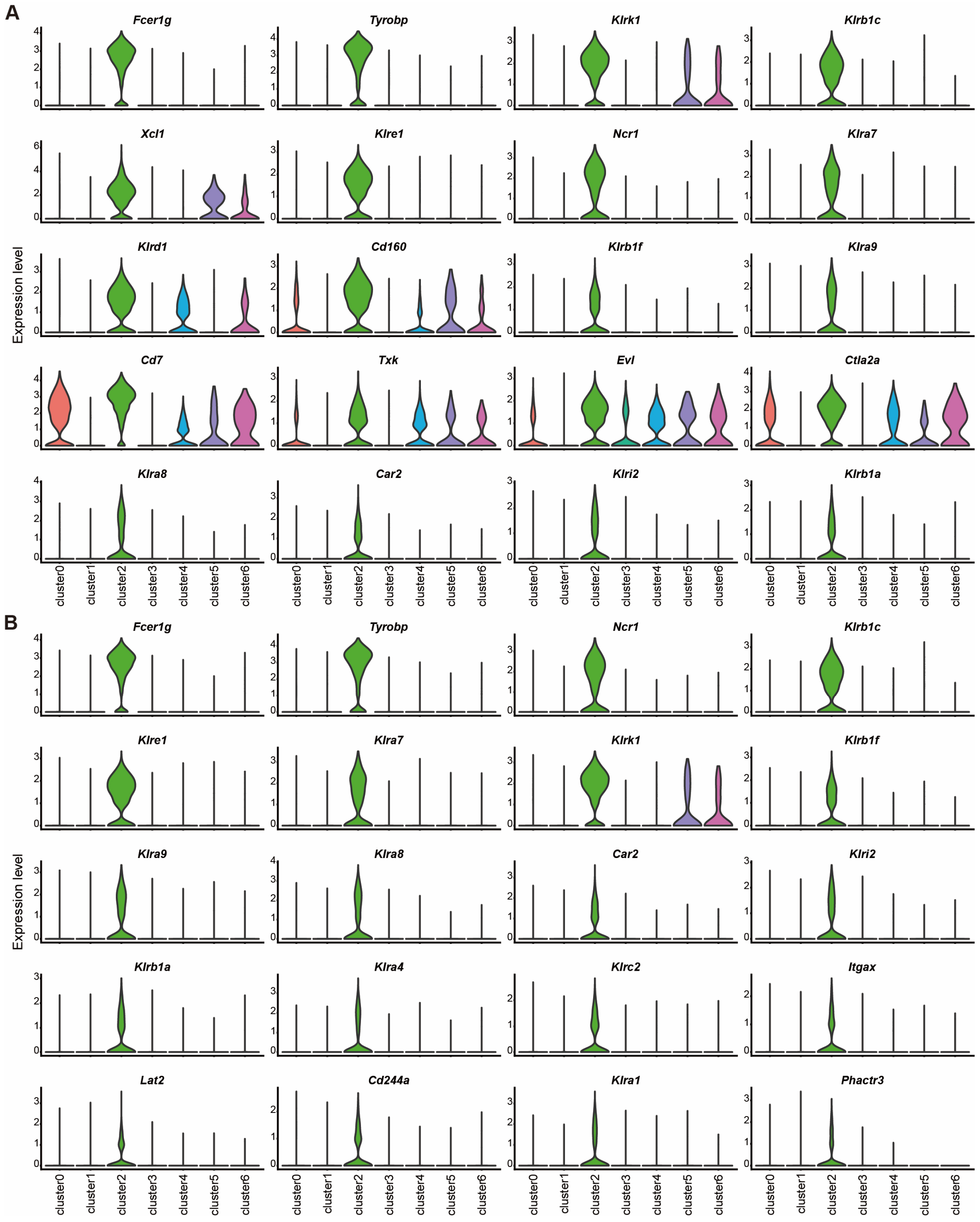

**Figure S7. The expression distribution of differentially expressed genes for the cluster 2 cell population in the scRNA-seq dataset from mouse models of multiple myeloma.** (**A**) The violin plots show the expression distribution of the top 20 differentially expressed genes identified by the Wilcoxon rank-sum test across different cell populations. Eight of these genes lack specificity for cluster 2 cell population. (**B**) The violin plots show the expression distribution of the top 20 differentially expressed genes identified by CSFeatures across different cell populations. Nineteen of these genes exhibit strong cell type-specific expression.

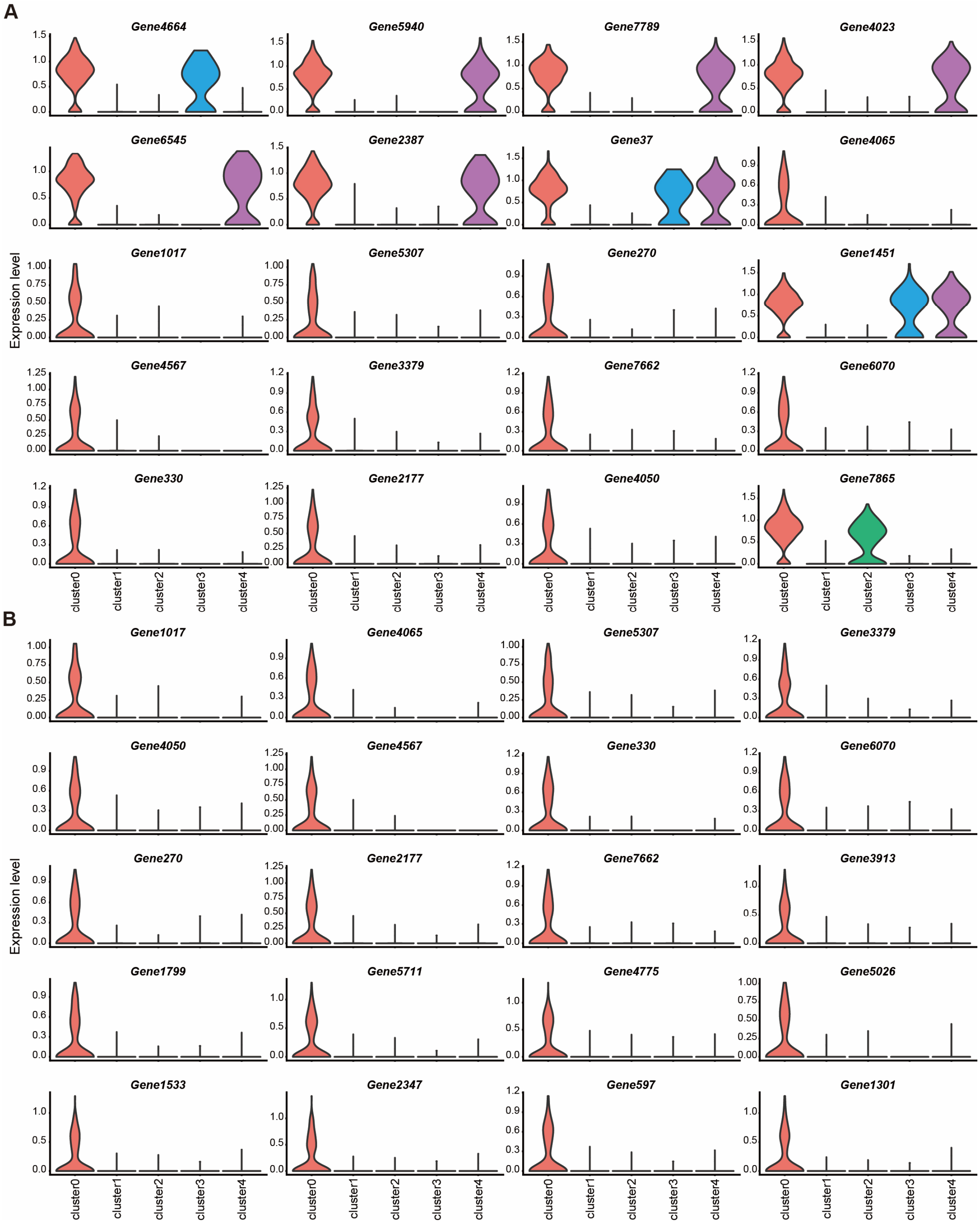

**Figure S8. The expression distribution of differentially expressed genes for the cluster 0 cell population in the simulated scRNA-seq dataset.** (**A**) The violin plots show the expression distribution of the top 20 differentially expressed genes identified by the Wilcoxon rank-sum test across different cell populations. Nine of these genes lack specificity for cluster 0 cell population. (**B**) The violin plots show the expression distribution of the top 20 differentially expressed genes identified by CSFeatures across different cell populations. All genes exhibit strong cell type-specific expression.

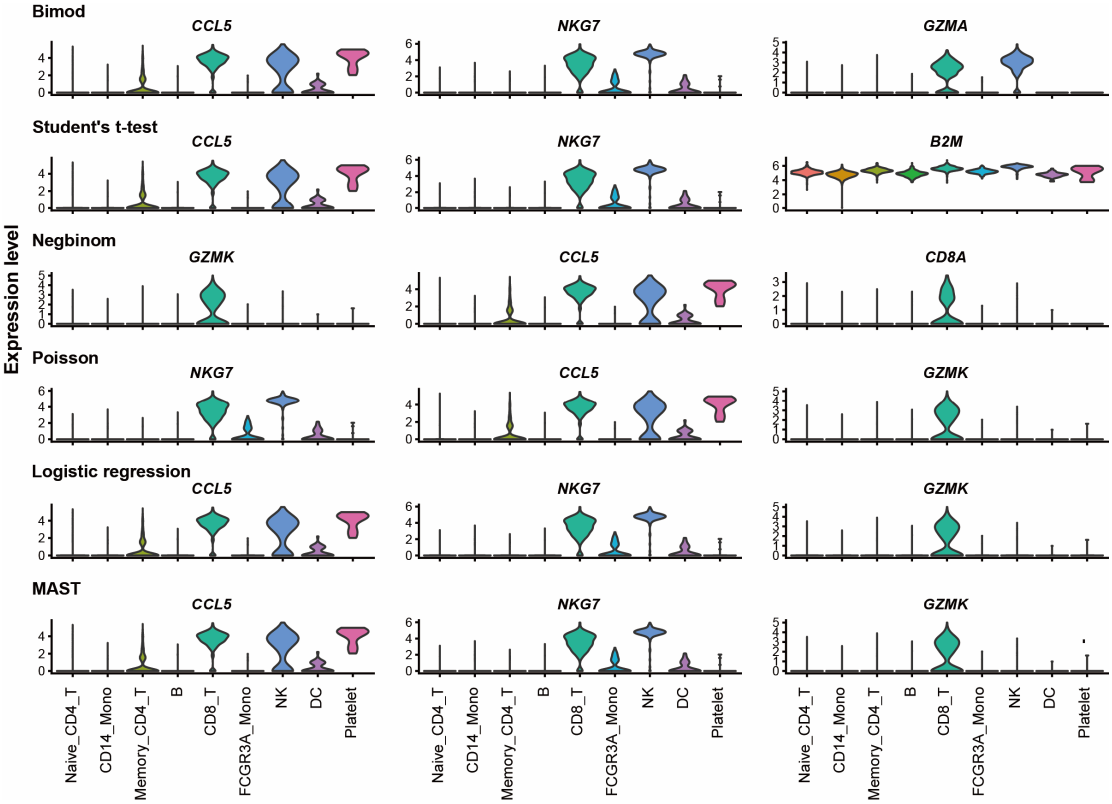

**Figure S9. Performance comparison of various differential analysis methods.** Using the top three differentially expressed genes for CD8 T cells in the PBMC dataset as an example, each row represents the results of a different differential analysis method. From top to bottom, the methods are bimod, Student's t-test, negbinom, Poisson, logistic regression, MAST.

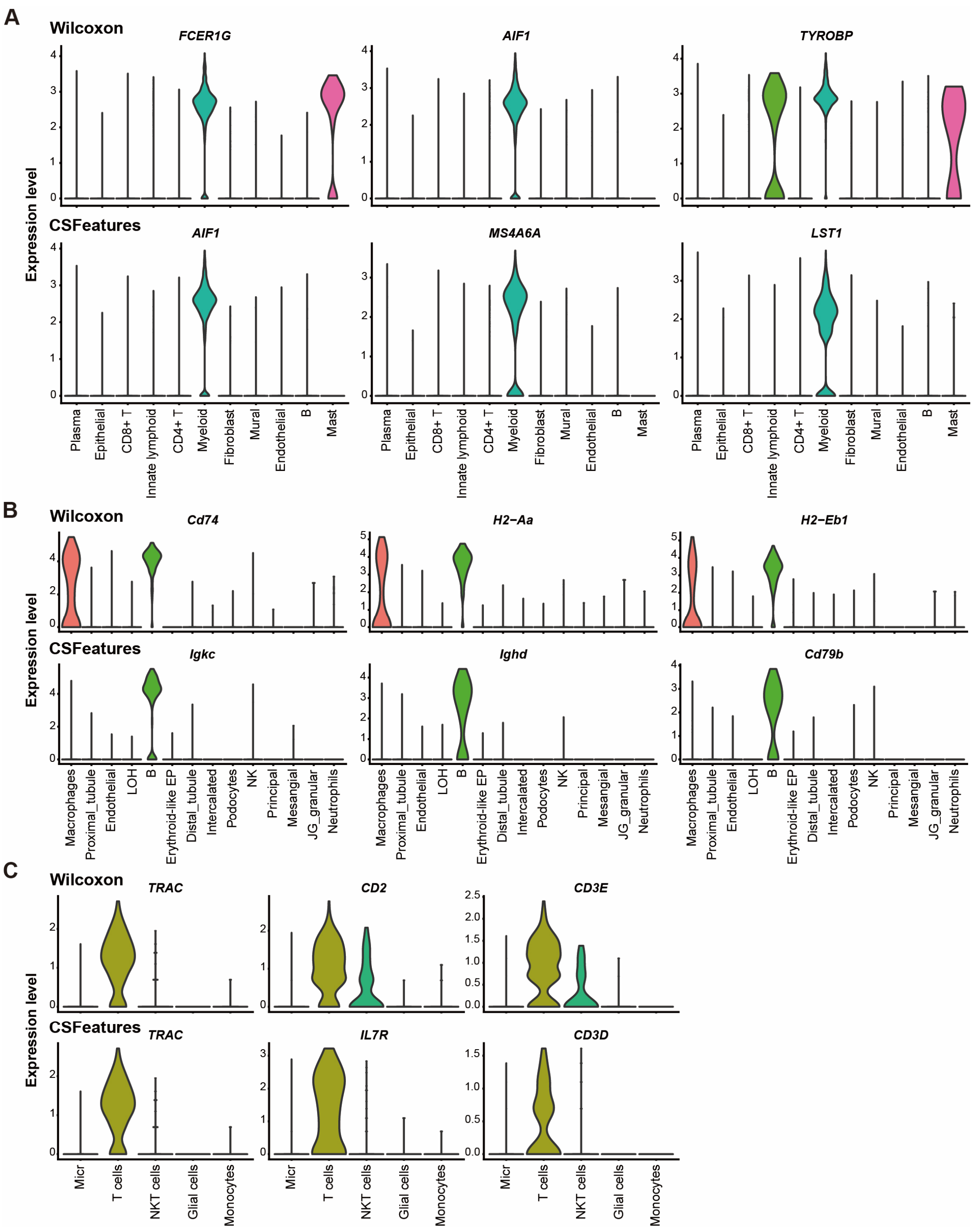

**Figure S10. Comparison of CSFeatures and the Wilcoxon rank-sum test in identifying differentially expressed genes (DEGs) across scRNA-seq datasets generated by various sequencing technologies.** (**A**) Expression distribution of the top three DEGs for myeloid cell population in the HCC dataset generated by 10x Chromium, identified by the Wilcoxon rank-sum test (top) and CSFeatures (bottom). (**B**) Expression distribution of the top three DEGs for B cell population in the mouse kidney dataset generated by Drop-seq, identified by the Wilcoxon rank-sum test (top) and CSFeatures (bottom). (**C**) Expression distribution of the top three DEGs for T cell population in the human glioblastoma dataset generated by CEL-seq2, identified by the Wilcoxon rank-sum test (top) and CSFeatures (bottom).

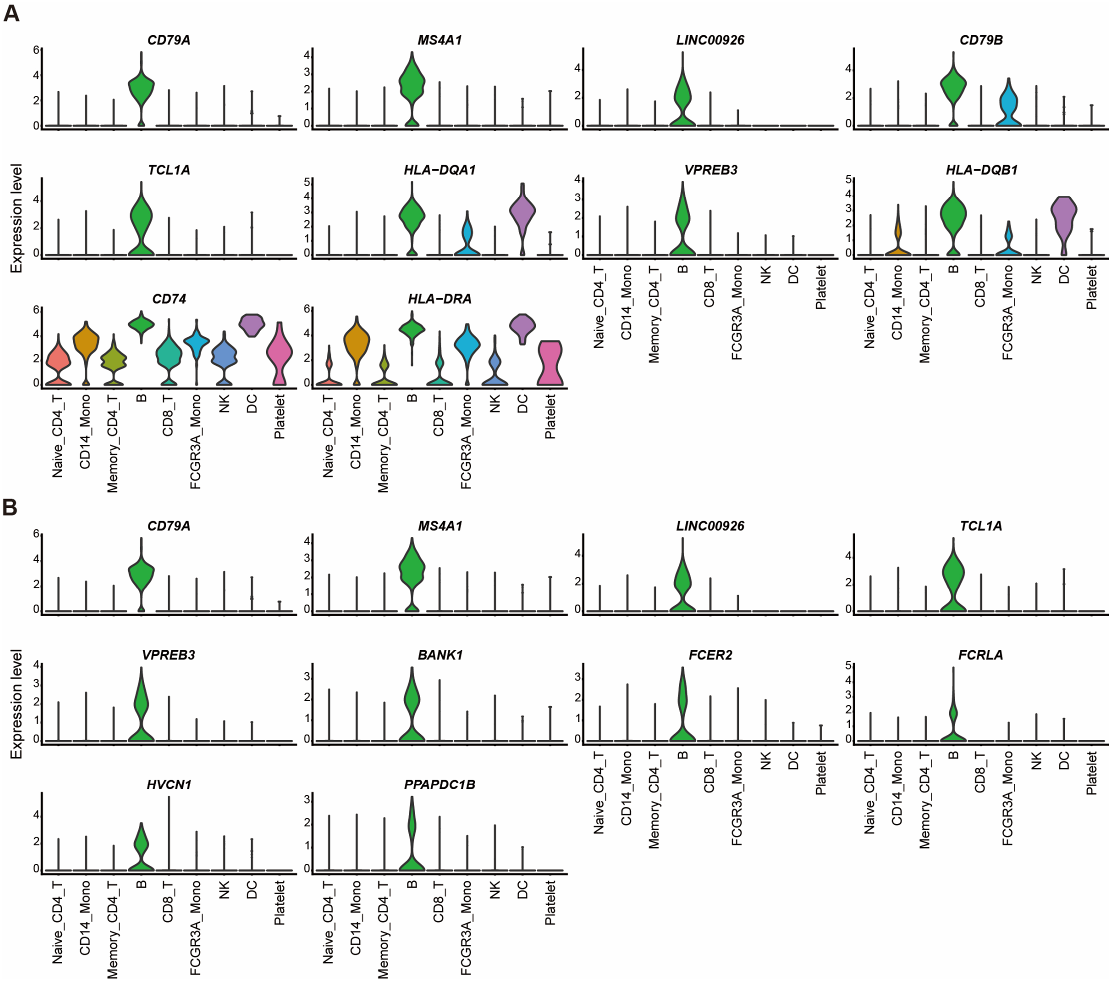

**Figure S11. The expression distribution of differentially expressed genes for the B cell population in the PBMC dataset.** (**A**) The violin plots show the expression distribution of the top 10 differentially expressed genes identified by the Wilcoxon rank-sum test across different cell populations. Half of these genes lack specificity for B cells. (**B**) The violin plots show the expression distribution of the top 10 differentially expressed genes identified by CSFeatures across different cell populations. All genes exhibit strong B cell type-specific expression.

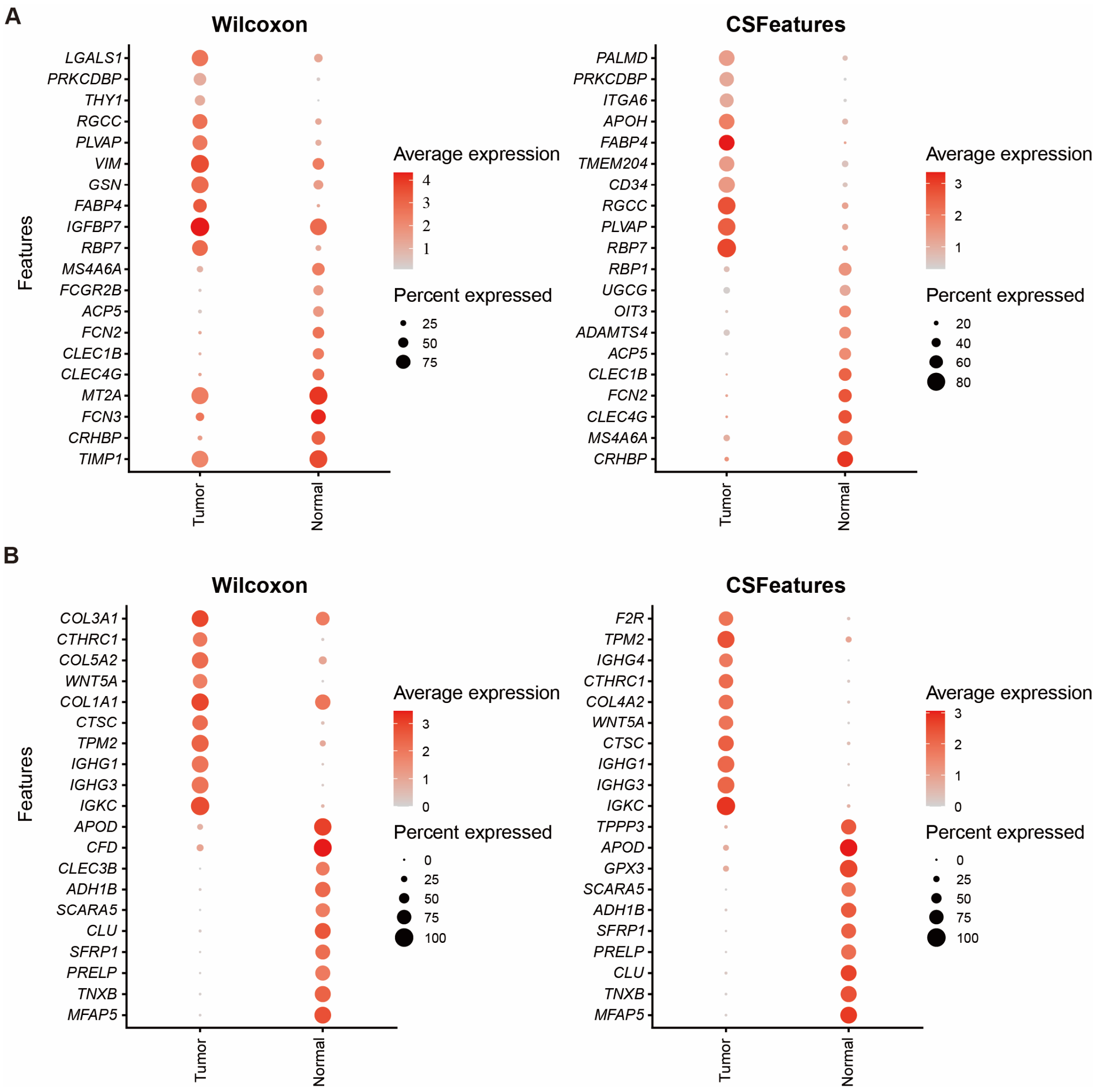

**Figure S12. Expression distribution of differentially expressed genes across varying cell population states.** (**A**) For the endothelial cell population in the lung cancer dataset, the top 10 genes with the most significant upregulation or downregulation in tumor tissue were identified by the Wilcoxon rank-sum test (left) and CSFeatures (right). The color intensity reflects the gene expression levels, while bubble size represents the proportion of cells expressing the gene. (**B**) For the fibroblast population in the gastric cancer dataset, the top 10 genes with the most significant upregulation or downregulation in tumor tissue were identified by the Wilcoxon rank-sum test (left) and CSFeatures method (right). The color intensity reflects the gene expression levels, while bubble size represents the proportion of cells expressing the gene.

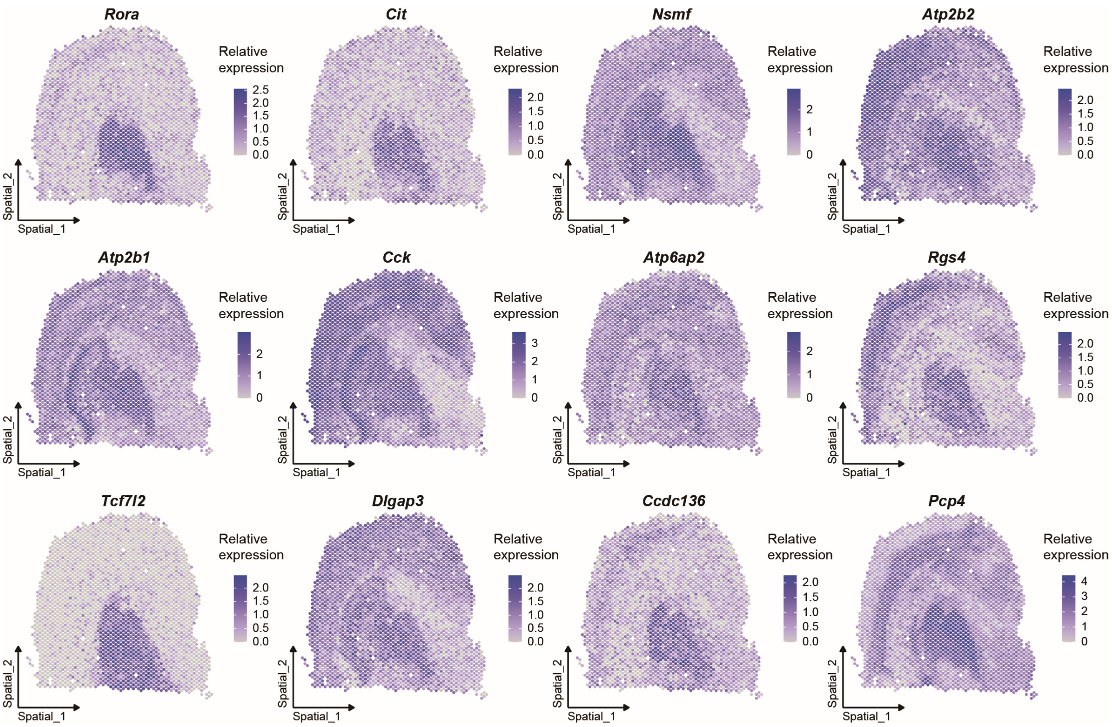

**Figure S13. The expression distribution of differentially expressed genes identified by SpaGCN for Thalamus_1 population in the mouse brain spatial transcriptomics dataset.** The top 12 differentially expressed genes identified by SpaGCN in the Thalamus_1 population exhibit poor cell type specificity. All genes show high expression not only in the Thalamus_1 population but also broadly in other cell populations. The color intensity reflects the gene expression levels.

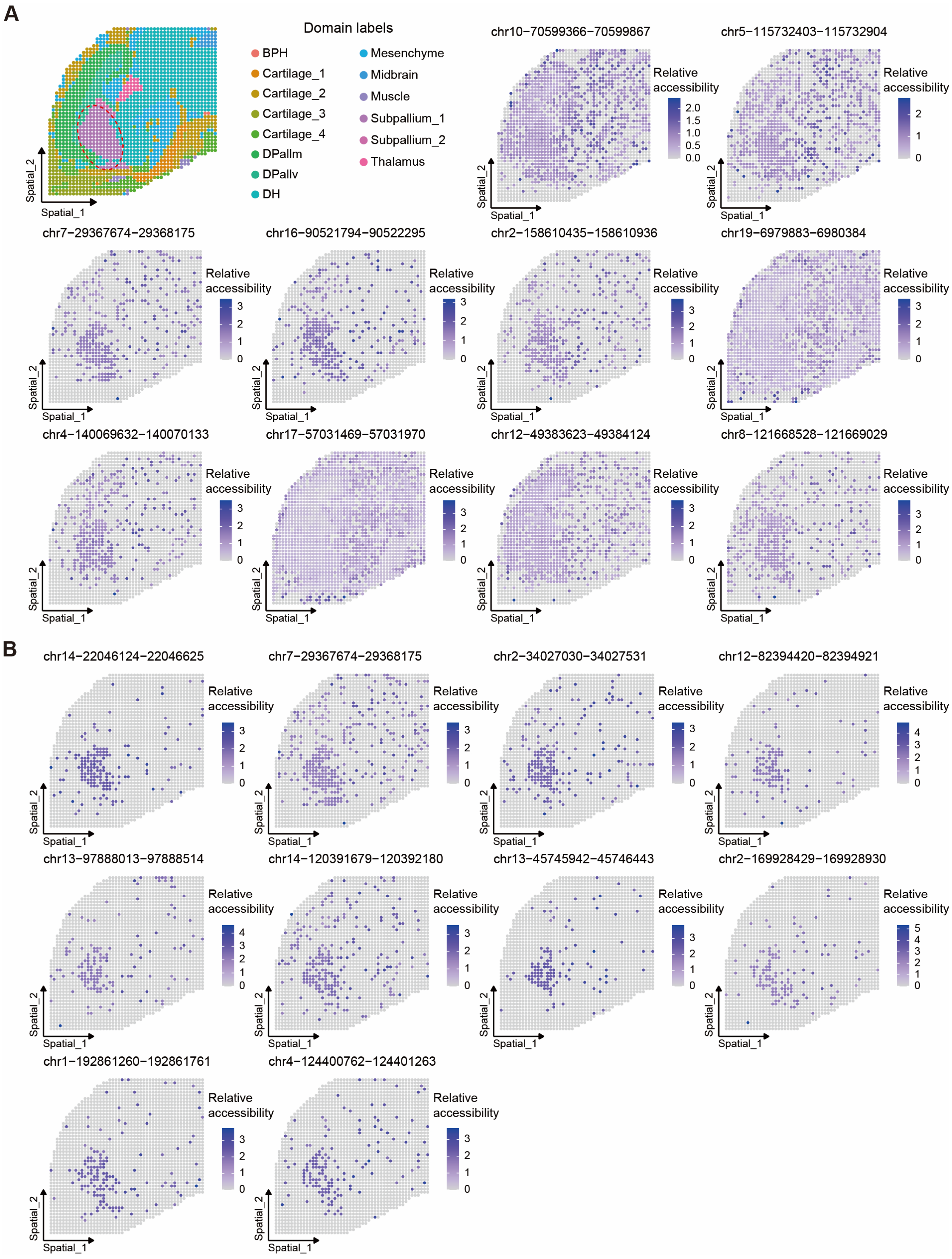

**Figure S14. CSFeatures identifies differentially accessible regions with spatial patterns.** In the mouse embryo spaATAC-seq dataset, we used the Subpallium_1 population as an example. (**A**) The top 10 differentially accessible regions identified by LR. All regions are highly accessible in the Subpallium_1 population and other cell populations. The color intensity reflects the chromatin accessibility levels. (**B**) The top 10 differentially accessible regions identified by CSFeatures. The vast majority of regions demonstrate specificity for the Subpallium_1. The color intensity reflects the chromatin accessibility levels.

### Supplementary Tables

**Table S1.** The top 10 unique DEGs for fibroblasts ranked by CSFeatures and their functional links.

| Genes | Descriptions | References |
|---|---|---|
| *PDGFRA* | Cardiac fibroblast survival and proliferation require *PDGFRA* signaling. | Ivey et al.(4) |
| *PDPN* | The gene *PDPN* is a marker of fibroblast. | Friedman et al.(5) |
| *DIO2* | The gene *DIO2* is a cytokine secreted by synovial fibroblasts. | Pörings et al.(6) |
| *ISLR* | The gene *ISLR* is a marker of fibroblast. | Takahashi et al.(7) |
| *CXCL14* | The gene *CXCL14* is a marker of fibroblast. | Li et al.(8) |
| *EFEMP1* | The gene regulates fibroblast activity. | Cosentino et al.(9) |
| *FBLN5* | The gene *FBLN5* accelerates elastic fibre assembly in human skin fibroblasts. | Katsuta et al.(10) |
| *FENDRR* | The gene *FENDRR* affects lung fibroblast proliferation. | Senavirathna et al.(11) |
| *MMP23B* | The gene *MMP23B* is a matrix metalloproteinase produced by fibroblasts. | Podstawski et al.(12) |
| *PID1* | The gene *PID1* affects fibroblast cell cycle. | Monteleone et al.(13) |

**Table S2.** The top 10 unique DEGs for fibroblasts ranked by Wilcoxon rank-sum test and their functional links.

| Genes | Descriptions | References |
|---|---|---|
| *CALD1* | The gene *CALD1* is a marker of fibroblast. | Wu et al.(14) |
| *C1R* | None | None |
| *COL1A1* | The gene *COL1A1* is a marker of fibroblast. | Li et al.(15) |
| *COL6A1* | None | None |
| *COL6A2* | None | None |
| *COL1A2* | The gene *COL1A2* is a marker of fibroblast. | Hutchenreuther et al.(16) |
| *MMP2* | The gene *MMP2* is a matrix metalloproteinase produced by fibroblasts. | Le et al.(17) |
| *FSTL1* | The gene *FSTL1* is the fibroblast-derived growth factor. | Loh et al.(18) |
| *TPM1* | None | None |
| *PCOLCE* | None | None |

**Table S3.** Summary of the datasets used in this study.

| Category | Datasets | No. of cells | No. of features | Technology |
|---|---|---|---|---|
| scRNA | Human PBMC | 2,700 | 13,714 | 10X Chromium |
| | Human gastric cancer | 11,368 | 25,255 | 10X Chromium |
| | Human hepatocellular carcinoma | 71,915 | 25,712 | 10X Chromium |
| | Human leukemia | 11297 | 22303 | 10X Chromium |
| | Mouse multiple myeloma | 12978 | 20350 | 10X Chromium |
| | Mouse kidney | 16,015 | 23,511 | Drop-Seq |
| | Human glioblastoma | 4,460 | 23,183 | CEL-Seq2 |
| | Simulated dataset | 2000 | 8000 | None |
| scATAC | Mouse forebrain | 2,088 | 162,398 | Preissl et al.(19) |
| | Mixture of human cell lines | 549 | 157,358 | Liu et al.(20) |
| ST | Mouse brain | 2,686 | 32,245 | 10X Visium |
| spaATAC | Mouse embryo brain | 2,129 | 37,475 | 10X Genomics |
